# Synaptic and neural pathway redundancy enables the robustness of a sensory-motor reflex and promotes predation escape in *C. elegans*

**DOI:** 10.1101/2025.09.18.677212

**Authors:** Haoming He, Eugenia King Hin Fong, Sandeep Kumar, Ho Ming Terence Lee, Andrew M. Leifer, Martin Chalfie, Chaogu Zheng

## Abstract

As a basic unit of the nervous system, the sensory-motor reflex circuit is fast and robust. However, it is not entirely clear how this robustness is achieved, given that various genetic perturbations can disrupt the function of the sensory neurons. By mapping the molecular basis of neuronal connections in the touch response circuit of *Caenorhabditis elegans*, we found prevalent genetic redundancy at neural pathway, synaptic, and molecular levels, which ensures that sensory signals can be relayed to command interneurons that control motor output. We also discovered developmental remodeling of the anterior circuit, which leads to the pruning of larval synapses, establishment of a second pathway that activates additional interneurons, and lateralization of the circuit. Finally, we found that the synapses that appeared to be functionally redundant in a simple touch assay contribute to the extent of reversal response in an additive manner, which may help the organism escape from predators.

## INTRODUCTION

Sensory-motor reflex circuits are evolutionarily conserved building blocks for complex nervous system in animals. The circuit consists of sensory neurons, interneurons, and motor neurons.^1^ The sensory neurons, either directly or indirectly through a sensory organ, detect environmental stimuli (such as light, sound, smell, touch, and temperature) and transmit them into electrical signals. The signal travels through neural pathways to reach interneurons, activating or inhibiting them through synapses. Interneurons then control the activity of motor neurons, which in turn activate the effector muscles to evoke motor responses. Sensory neurons can also be directly connected to motor neurons in monosynaptic reflexes.^2^ The sensory-motor reflexes are known to be quick and robust and play important roles in helping the host organism avoid danger. However, a paradox for this robustness is that sensory neurons are often vulnerable to genetic mutations. For example, genetic screens searching for mutants with defects in sensory perception (e.g., phototaxis in Drosophila^3^, chemotaxis^4^ and mechanosensation^5^ in *C. elegans*, acoustic startle response in zebrafish^6^ and mice^7^) often lead to the discovery of genes involved in the development and function of the sensors but rarely uncover genes involved in synaptic transmission. This fact suggests that the robust response of the reflex may rely on efficient and potentially redundant signal transmission downstream of the sensory neurons. In many cases, the molecular and cellular basis of such robustness and redundancy is unclear.

In this study, we focus on the gentle touch reflex in the nematode *Caenorhabditis elegans*. Initial work mapped both the physical and functional connectivity of the touch circuit, which is comprised of six mechanosensory neurons (known as the touch receptor neurons or TRNs), several interneurons, and three types of motor neurons.^8,9^ Among the six TRNs, ALML, ALMR, PLML, and PLMR are embryonic neurons, and AVM and PVM are postembryonic neurons. The anterior TRN subtype ALM neurons are connected through gap junctions with the interneuron AVD that controls backward movement and through inhibitory chemical synapses with the interneuron PVC that controls forward movement, which allows anterior touch to induce backward movement and inhibit forward movement. The posterior TRN subtype PLM neurons do the opposite, so that tail touches induce forward movement and prevent backward movement (Figure 1A). At the late larval stages, the anterior circuit is remodeled upon the joining of AVM, which not only connects with ALML and ALMR *via* gap junctions to allow lateral integration but also serves as a signaling hub to activate AVD through gap junctions. Moreover, AVM generates motor output through a secondary pathway that involves AVB and AVA interneurons in adults (Figure 1A). AVM makes presumably inhibitory chemical synapses with AVB, which is then connected to AVA that controls backward movement.^8,10^ This connectivity suggests that in adults anterior touch may promote backward movement through two independent and potentially redundant neural pathways. Indeed, ablating either AVD or AVA did not result in anterior touch sensitivity but ablating both did,^8^ confirming the redundancy between the two interneuron pathways.

**Figure 1.**
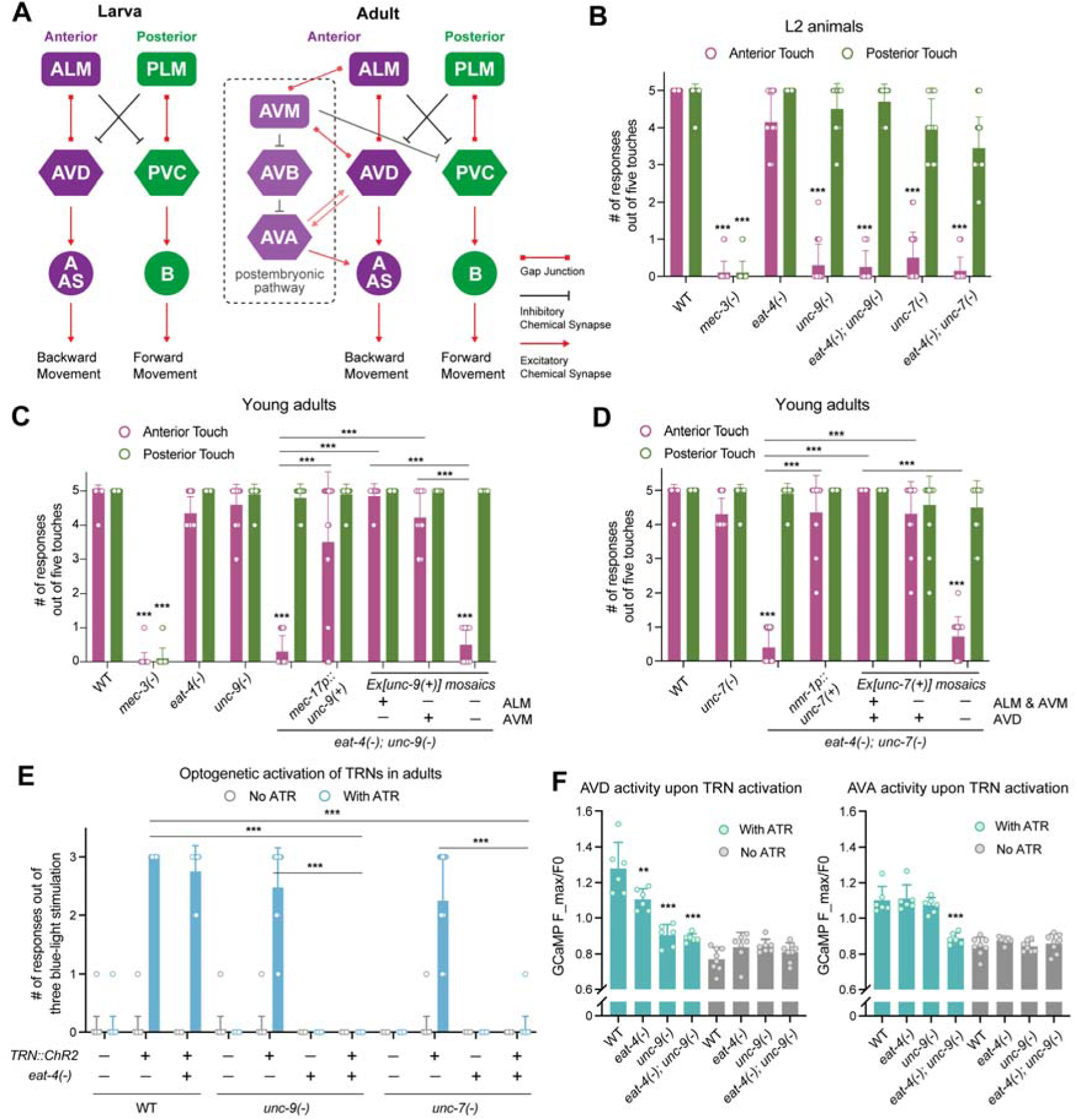
UNC-9::UNC-7 gap junctions and glutamatergic synapses redundantly regulate anterior touch response in adults. (A) Connectivity in the anterior touch circuit in young larvae and adults. (B) The number of responses out of five anterior or posterior touches in the indicated strains; *eat-4(ky5)*, *unc-9(e101)*, and *unc-7(e5)* alleles were used. *mec-3(u184)* mutants had defective TRN differentiation and were used as a negative control. (C-D) Touch responses of the indicated strains. For *unc-9* mosaic analysis, *unkEx459[unc-9(+); mec-17p::TagRFP; nmr-1p::GFP]* was used; for *unc-7* mosaic analysis, *unkEx629[unc-7(+); mec-17p::TagRFP; nmr-1p::GFP]* was used. Since *nmr-1p::GFP* labels multiple interneurons in the head, for “AVD-“ we selected animals with no GFP signal in the head. For (B-D), N = 20 for all except for the mosaic cases where N = 10∼15. Triple asterisks indicate *p* < 0.001 in a post-ANOVA Tukey’s test comparing the mutant with wild-type or selected pairs. (E) The number of reversal responses out of three blue light stimulations of the head of animals carrying the *uIs94[mec-4p::ChR2::YFP]* transgene in the presence or absence of *all-trans*-retinal (ATR). N = 20. (F) Calcium signal change in AVD and AVA upon activation of TRNs in animals carrying the *wtfIs46[mec-4p::Chrimson]* and *sraIs49[nmr-1p::GCamp3]* transgenes and a *lite-1(lf)* mutation. The ratio of maximum cell body GCaMP intensity after the stimulation to the intensity before the stimulation is shown. N ≥ 6. Two and three asterisks indicate *p* < 0.01 and 0.001, respectively, in a post-ANOVA Dunnett’s test in comparison with the wild-type animals. Mean ± SD is shown in this Figure.

Despite the known physical map and functional connectome of the touch circuit from 40 years ago, the molecular basis of the neuronal connections in the circuit is largely unknown. Saturated forward genetic screens searching for touch-insensitive mutants led to the discovery of 18 genes required for the touch reflex response.^5,11^ Strikingly, all of them regulate either the differentiation or mechanosensory functions of the TRNs and none regulates the synaptic transmission downstream of the sensory neurons. Later, an RNAi screen of all the essential genes (whose mutation cannot be isolated from the mutagenesis screen) led to the identification of 61 more genes involved in touch sensation, but few or none of them specifically regulates synapses.^12^ These results strongly hinted at genetic redundancy governing the activation of downstream interneurons.

Here, through a candidate approach, we identified the synaptic proteins that form the molecular basis of the neuronal connection in the touch circuit and discovered layers of redundancy achieved through distinct mechanisms at the gene, synapse, and neural pathway levels. We also observed synaptic pruning during developmental remodeling of the circuit that helps build the redundant interneuron pathways. Finally, although the redundant genes, synapses, and pathways ensure robust touch reflex in adult animals, they all contribute to the extent of the touch response in an additive manner and enable the escape from predation by carnivorous nematodes. Overall, through comprehensive genetic studies, our work provides an example to illustrate how redundancy is built at the neuronal connections in a sensory-motor reflex to ensure its robustness.

## RESULTS

### Molecular basis of the TRN-interneuron connections in the anterior touch circuit

Since the touch avoidance response is thought to be driven by the excitatory gap junctions between the mechanosensory TRNs and the interneurons,^8^ we first sought to identify the molecular basis of these electrical synapses in the circuit. In invertebrates, innexin proteins form hexameric or octameric hemichannels that connect across the intercellular space to form gap junctions between two neurons.^13,14^ The *C. elegans* genome contains 25 innexin genes,^15^ and nervous system-wide expression mapping using fosmid-based reporters identified the innexins expressed in the touch circuit^16^. Many of these reported expressions were confirmed by single-cell transcriptomic data^17^, although there are some discrepancies between the two datasets (Table S1). In total, ten innexins showed expression in the circuit and may be involved in constructing the gap junctions. Since the ALM::AVD gap junction enables backward movement upon anterior touch in young larva, we tested the mutants of all innexin genes expressed in ALM and AVD neurons (Table S2) and found that mutations in *unc-9* and *unc-7* caused defects in anterior touch response in young larva (L2-L3 animals), suggesting that the ALM::AVD gap junction depends on UNC-9 and UNC-7 (Figure 1B and S1A).

Interestingly, both *unc-9(-)* and *unc-7(-)* animals regained anterior touch sensitivity in L4 and adults (Figure 1C, 1D, and S1A) likely because of the incorporation of AVM into the anterior circuit and the establishment of the AVM-AVB-AVA pathway in late larval development. To confirm this idea, we disabled the chemical synapses from the glutamatergic TRNs^18^ by removing the vesicular glutamate transporter *eat-4*/*vglut*, which blocks the connection between AVM and AVB. We found that although *eat-4(-)* single mutants were touch-sensitive, both *unc-9(-); eat-4(-)* and *unc-7(-); eat-4(-)* double mutants were touch-insensitive at the anterior (Figure 1C and 1D), suggesting that the gap junctions and chemical synapses serve redundant functions in mediating anterior touch response in adults. As a control, these double mutants were still touch-sensitive at the posterior, indicating normal mechanotransduction in the TRNs but defective synaptic transmission in the anterior circuit.

UNC-9 is known to form gap junction with UNC-7 with specific directionality from UNC-9 to UNC-7 hemichannels.^19,20^ Moreover, according to the fosmid reporter expression pattern, *unc-7* is not expressed in the TRNs of well-fed adults.^16^ Thus, we hypothesized that UNC-9 in the anterior TRNs may form intercellular gap junctions with UNC-7 in the anterior interneuron AVD. Supporting this hypothesis, expression of the wild-type *unc-9(+)* from a TRN-specific *mec-17* promoter rescued anterior touch sensitivity in *unc-9(-) eat-4(-)* double mutants (Figure 1C). More importantly, a mosaic analysis found that *unc-9(+)* expression in either ALM or AVM was able to rescue the anterior touch response, suggesting that both ALM::AVD and AVM::AVD gap junctions rely on UNC-9 (Figure 1C). For *unc-7*, the expression of a wild-type copy from an interneuron-specific *nmr-1* promoter was able to restore the anterior touch sensitivity in the *unc-7(-); eat-4(-)* mutants, and a mosaic analysis showed that *unc-7(+)* expression was not required in the ALM and AVM but was required in the AVD for the touch response (Figure 1D). These results indicated that the UNC-9::UNC-7 gap junction is essential for the connectivity between TRNs and AVD, while in adults this pathway is redundant due to the addition of the chemical synapse-dependent AVM-AVB-AVA pathway.

To bypass the need for mechanical stimulation, we optogenetically activated ALM and AVM by expressing channelrhodopsin in the TRNs and shining blue light to the head of the animal. We found that although *unc-9(-)*, *unc-7(-)*, and *eat-4(-)* single mutants were all able to respond to the head stimulation by reversal, the *unc-9(-); eat-4(-)* and *unc-7(-); eat-4(-)* double mutants did not respond to the blue light, suggesting that the circuit connection downstream of the TRNs was mediated redundantly by UNC-9::UNC-7 gap junctions (ALM/AVM::AVD) and glutamatergic chemical synapses (AVM-AVB) (Figure 1E). To test the contribution of AVM to the circuit, we ablated AVM using a UV laser and found that the animal could still respond to optogenetic stimulation in the head, suggesting that the ALM::AVD connection is sufficient to drive the touch avoidance response in both larvae and adults. As expected, ablating AVM in the *unc-9(-)* mutants eliminated the reversal response (Figure S1B), since the ALM::AVD gap junction requires *unc-9*. These results further confirm the above model of synaptic connection in the touch circuit.

Moreover, by expressing the red-light activated Chrimson in the TRNs and a GCaMP protein in the interneurons, we recorded the activities of AVD and AVA interneurons (both promoting backward movement) upon the activation of ALM and AVM in the anterior of the animal. We found that the activation of AVD depended on UNC-9 as expected, although we also observed a slightly reduced AVD activation in *eat-4(-)* mutants, likely because AVA contributes to AVD activation (Figure 1F). However, since the *eat-4(-)* mutants were touch-sensitive, the small reduction in AVD activity did not significantly impact behavioral response in a touch assay. Interestingly, *eat-4(-)* mutants did not show reduced AVA activity, suggesting that AVA can also be activated through the connection with AVD, which is consistent with previous findings.^21,22^ Importantly, anterior TRN-triggered AVA activation was completely blocked in *eat-4(-); unc-9(-)* double mutants, confirming that both ALM/AVM::AVD and AVM-AVB pathways can redundantly activate AVA.

Furthermore, we conducted a forward genetic screen to identify genes involved in the ALM/AVM::AVD pathway using *eat-4(-)* mutants as the starter strain and searched for mutants that were touch-insensitive at the anterior but sensitive at the posterior. After screening ∼5,000 haploid genomes, we identified two mutants. One was a *lf* allele of *unc-42*, which regulates the specification of interneuron cell fate in AVD;^23^ the other was a nonsense mutation (Q334*) in *unc-9* (Figure S1C). Thus, an unbiased approach independently identified UNC-9 as a critical component of the ALM/AVM::AVD connection.

### INX-1::INX-1 and UNC-9::UNC-7 gap junctions act redundantly for the PLM::PVC connection in the posterior touch circuit

When analyzing the mutants of the ten innexins expressed in the circuit, we found that none of the single mutants disrupted posterior touch sensitivity, indicating potential genetic redundancy (Table S2). Given that *unc-9* was involved in the anterior circuit, we constructed double mutants combining *unc-9(-)* with the *lf* alleles of other innexins and found that only *inx-1(-) unc-9(-)* double mutants showed defects in posterior touch response (Figure 2A and Table S2). Both a nonsense allele *(gk580946)* and the deletion allele (*unk29*; Figure S2A) of *inx-1* produced similar phenotypes in combination with the *unc-9(e101)* mutant; *unk29* was used as *inx-1(-)* in subsequent studies.

**Figure 2.**
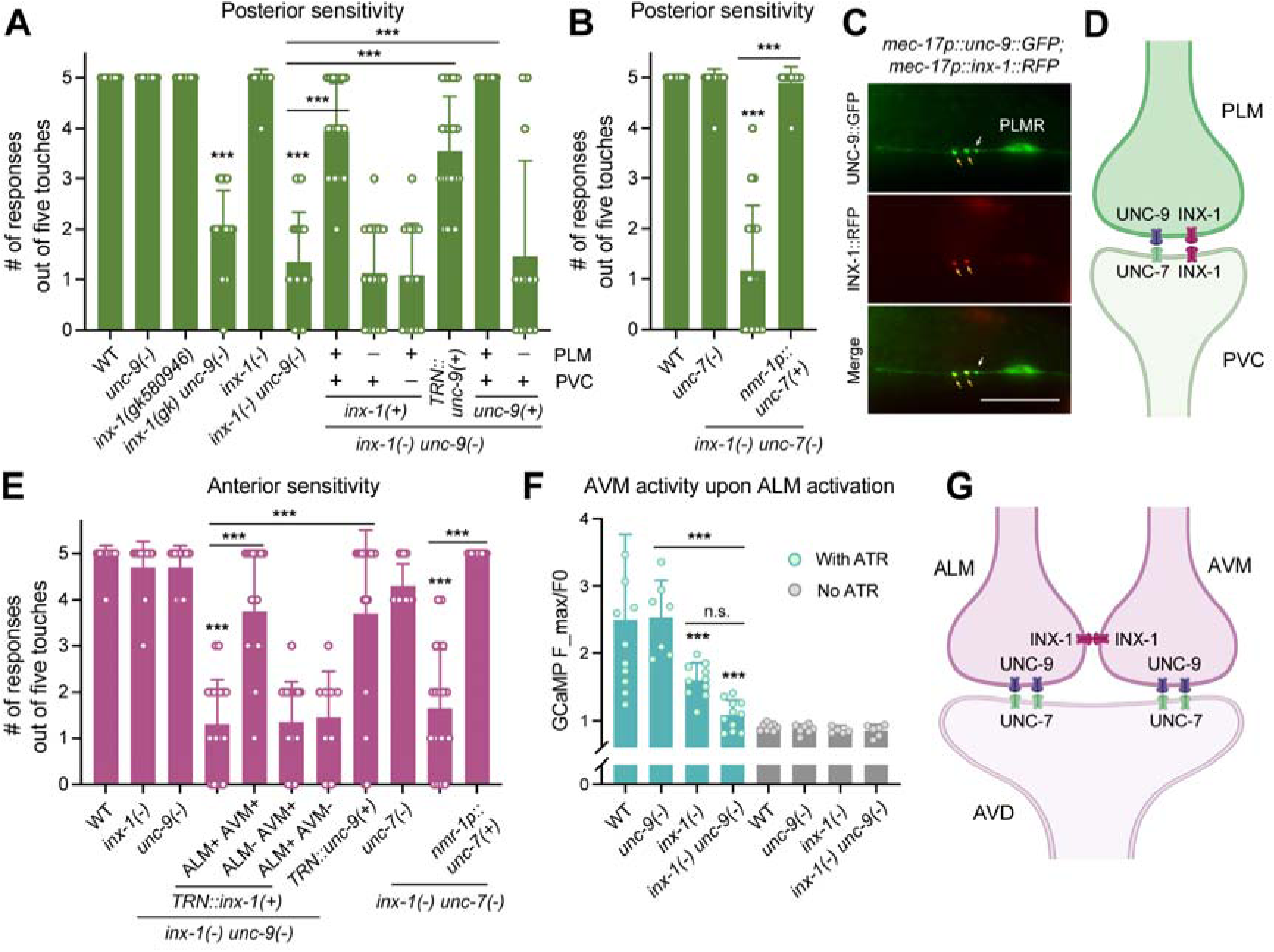
UNC-9 and INX-1 redundantly regulate touch sensitivity in both anterior and posterior sides. (A-B) The number of responses out of five posterior touches in various strains; *inx-1(gk580946 or gk)*, *inx-1(unk29 or -)*, *unc-9(e101)*, and *unc-7(e5)* alleles were used. For *inx-1* mosaics, *unkEx652[mec-17p::inx-1a; mec-17p::TagRFP; nmr-1p::inx-1a; nmr-1p::GFP]* were used; for *unc-9* mosaics, *unkEx459[unc-9(+); mec-17p::TagRFP; nmr-1p::GFP]* was used. N = 20. Triple asterisks indicate *p* < 0.001 in a Tukey’s multiple comparison test. (C) Representative images of UNC-9::GFP and INX-1::RFP puncta in the PLM anterior neurite. Orange arrows indicate the puncta with overlapping GFP and RFP signal, while the white arrows indicate GFP only puncta. Scale bars = 10 μm. (D) A schematic cartoon for the gap junctions formed between PLM and PVC. (E) The number of responses out of five anterior touches in various strains. For *inx-1* mosaics, *unkEx643[mec-17p::inx-1a; mec-17p::TagRFP]* was used. N = 20. (F) Calcium signal change in AVM upon optogenetic activation of ALMR in animals carrying the *unkEx902[T20B3.14p::Chrimson; mec-17p::GCaMP6s]* transgene. *T20B3.14* promoter is expressed in the ALM but no other TRNs. N = 7 for unc-9(e101) and N ≥ 10 for others. (F) A schematic cartoon for the gap junctions formed among ALM, AVM, and AVD in the anterior touch circuit. Mean ± SD is shown in this Figure.

To understand where *inx-1* and *unc-9* function in the PLM::PVC connection that mediates the posterior touch response, we first conducted a mosaic analysis for *inx-1* and found that only when *inx-1(+)* expression was restored in both PLM and PVC could the posterior sensitivity be restored in the *inx-1(-) unc-9(-)* double mutants (Figure 2A). This result suggests that *inx-1* is required in both pre- and post-synaptic cells, likely because INX-1 forms homotypic INX-1::INX-1 gap junctions.^24,25^ In contrast, TRN-only expression of *unc-9(+)* could rescue the posterior touch response in the double mutants, and a mosaic analysis found that *unc-9* was only required in the PLM neuron for the posterior sensitivity (Figure 2A). Thus, we speculated that UNC-9 in the PLM may pair with UNC-7 in the PVC to form the gap junctions.

Indeed, although *unc-7(-)* single mutants were sensitive in the tail, *inx-1(-) unc-7(-)* double mutants showed posterior insensitivity similar to the *inx-1(-) unc-9(-)* animals (Figure 2B). Importantly, *nmr-1* promoter-driven interneuron-specific expression of *unc-7(+)* fully rescued the posterior insensitivity in *inx-1(-) unc-7(-)* double mutants, suggesting that *unc-7* functions in the interneuron PVC. These results led to a model that two distinct types of gap junctions, INX-1::INX-1 and UNC-9::UNC-7, function redundantly to mediate the PLM::PVC connection in both larvae (Figure S2B) and adults (Figure 2A).

Moreover, we found that UNC-9::GFP and INX-1::RFP expressed in the PLM formed puncta that overlap with each other in the area where PLM forms gap junctions with PVC. This result suggests that UNC-9 and INX-1-based hemichannels may be homogenously localized in the same synaptic region, despite that they form functionally independent channels (Figure 2C and 2D). Interestingly, on average for every PLM process, we could find two puncta co-labelled by both UNC-9 and INX-1 and one UNC-9::GFP punctum that did not have an overlapping INX-1::RFP signal (Figure S2C). This extra UNC-9 punctum was mostly found on the PLMR axon. Since PLML/R also makes gap junctions with the bilaterally symmetric LUAL/R and the right-side only PVR in the same area,^9^ we suspect that the PLMR::PVR gap junction might only involve UNC-9 but not INX-1.

### INX-1::INX-1 gap junction connects ALM and AVM and contributes to the anterior touch response in adults

When conducting the touch tests of the innexin double mutants, we noticed that the anterior touch sensitivities of both *inx-1(-) unc-9(-)* and *inx-1(-) unc-7(-)* double mutants were reduced compared to the wild type, whereas the *inx-1(-)* single mutants were touch-sensitive (Figure 2E). Given that *inx-1(-)* larvae were also sensitive in the head (Figure S2B), we reasoned that unlike *unc-9* and *unc-7*, *inx-1* was not involved in the ALM::AVD connection.

Given that ALM and AVM also form gap junctions to allow the consolidation of anterior sensory signals, which may enhance anterior sensitivity (Figure 1A), we hypothesize that INX-1::INX-1 homotypic gap junction may mediate the ALM::AVM connection. To test this hypothesis, we rescued *inx-1* expression in the TRNs and conducted mosaic analysis in the *inx-1(-) unc-9(-)* mutant background. We found that only when *inx-1(+)* was expressed in both ALM and AVM could the anterior touch sensitivity be restored (Figure 2E), confirming that INX-1 functions in both cells. Moreover, by expressing Chrimson in the ALM neurons (using the *T20B3.14* promoter)^26^ and recording GCaMP signal in AVM neurons upon optogenetic activation of ALM, we found that AVM activity induced by ALM activation was much reduced in *inx-1(-)* mutants, supporting that INX-1 connects ALM and AVM (Figure 2F). In addition, ALM activation-triggered AVM activity was not reduced in *unc-9(-)* mutants, suggesting that UNC-9 may not be involved in the ALM::AVM gap junction (Figure 2F and 2G).

Interestingly, the anterior touch defects in *inx-1(-) unc-9(-)* and *inx-1(-) unc-7(-)* double mutants could be rescued by TRN-specific expression of *unc-9(+)* and interneuron-specific expression of *unc-7(-)*, respectively (Figure 2E). This result suggests that when the ALM/AVM::AVD connection is intact, the coupling among ALML, ALMR and AVM through INX-1::INX-1 gap junction is less significant in evoking motor response, probably because the signal can be propagated through both AVD and AVB interneurons. However, when the TRN::AVD pathway is blocked and the signal can only be passed down through the AVM-AVB-AVA pathway, the electrical coupling among the anterior sensors becomes critical for the avoidance behavior. Similarly, this coupling is also important when the AVM-AVB connection is blocked, since we found that the *eat-4(-); inx-1(-)* double mutants were moderately touch-insensitive in the head but not in the tail (Figure S2D). This touch defect can be rescued by re-expressing *inx-1(+)* only in the ALM and AVM and not in AVD, supporting that *inx-1* is not involved in the AVM::AVD gap junction. The above data suggest that the functional redundancy between the two interneuron pathways (i.e., TRN::AVD and AVM-AVB-AVA) is dependent on efficient coupling and signal consolidation among the three anterior sensors.

### The AVM-AVB-AVA pathway relies on multiple glutamate-gated chloride channels

Given that optogenetic excitation of ALM/AVM leads to the activation of AVA (which promotes backward movement) (Figure 1F) and that AVB and AVA are known to mutually inhibit each other,^27^ we hypothesized that AVM likely makes inhibitory chemical synapses with AVB, and the activation of AVM may hyperpolarize AVB and thus relieve the inhibition on AVA, leading to its activation. All TRNs use glutamate as the only neurotransmitter; deleting the vesicular glutamate transporter *eat-4* blocks the AVM-AVB pathway and results in anterior touch insensitivity when the TRN::AVD pathway is blocked by *unc-9(-)* mutations (Figure 3A). This anterior insensitivity can be rescued by TRN-specific expression of *eat-4(+)*, as well as *eat-4(+)* expression from a promoter that is active in AVM but not the other TRN subtypes (Figure 3A). This result suggests that glutamate release from AVM is critical for inducing backward movement through the chemical synapse-dependent AVM-AVB-AVA pathway.

**Figure 3.**
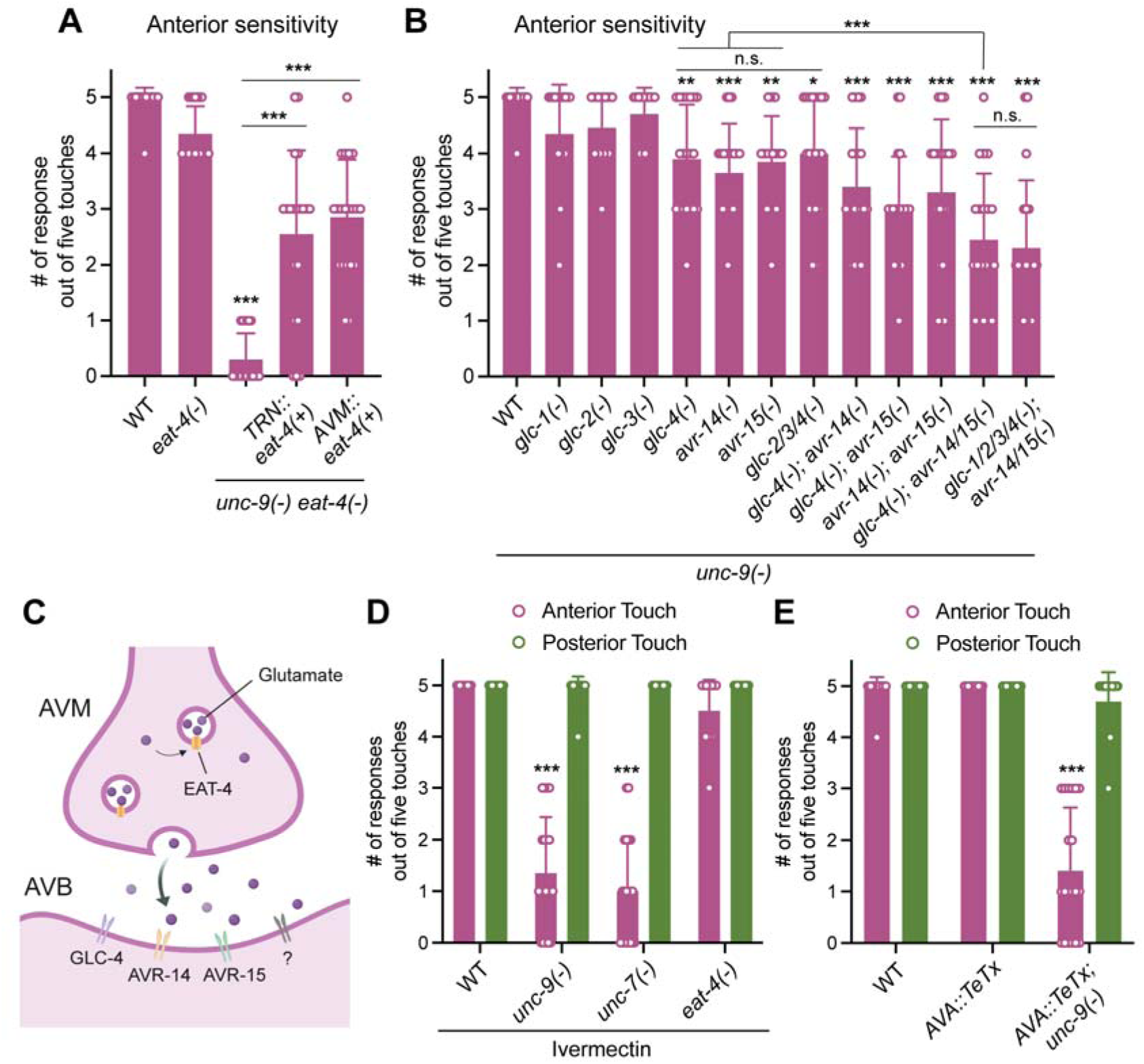
Glutamate-gated chloride channels mediate the AVM-AVB synapses and anterior touch response. (A) Anterior touch response in the *unc-9(e101); eat-4(ky5)* animals expressing *unkEx377[mec-17p::eat-4]* (for TRN-specific rescue) or *unkEx956[gcy-37p::eat-4]* (for AVM-specific rescue). N = 20. One, two, and three asterisks indicate *p* < 0.05, 0.01, and 0.001, respectively, in post-ANOVA Tukey’s tests. (B) Anterior touch response in the *unc-9(e101)* mutants carrying various combinations of mutations in the *glc* genes (see the key resources table for the strain information). (C) A schematic cartoon depicting the inhibitory glutamatergic synapses formed among AVM and AVB. For simplicity reasons, the GLC receptors were shown as homomeric, but they may also form heteromeric channels. “?” represents potentially unknown receptors that contribute to touch response. (D) The number of responses out of five anterior or posterior touches in animals treated with 5 nM ivermectin from embryonic stages. (E) Touch responses of animals carrying the *hpIs814[flp-18p::LoxP::eBFP::Stop::LoxP::TeTx::wCherry; twk-40p(short)::Cre]* transgene to express the neurotoxin in AVA neurons. Mean ± SD is shown in this Figure.

Based on the idea that AVM inhibits AVB, we next searched for inhibitory glutamate receptors, which are mostly the glutamate-gated chloride channels (GluCls). *C. elegans* genome contains at least six GluCl genes, namely *glc-1*, *glc-2*, *glc-3*, *glc-4*, *avr-14*, and *avr-15*.^28^ Single mutants of the GluCl genes did not affect touch sensitivity, while *glc-4(-)*, *avr-14(-)*, and *avr-15(-)* showed moderately reduced anterior sensitivity in double mutants with *unc-9(-)* (Figure 3B and S3). Previous single-cell transcriptome data confirmed that *glc-4* and *avr-14* are expressed in the AVB neurons.^17^ We then combined the GluCl mutants in the *unc-9(-)* background and found that the *glc-4(-) avr-15(-); avr-14(-); unc-9(-)* quadruple mutants had further reduced anterior touch response compared to the double mutants (Figure 3B), indicating that *glc-4*, *avr-14*, and *avr-15* function redundantly to promote touch-evoked reversal. Further removing *glc-1*, *glc-2*, and *glc-3* in the septuple mutants did not further decrease the sensitivity, suggesting that they may not be involved in the process (Figure 3B). Because the quadruple and even the septuple mutants, in which all GluCls were knocked out, were not as touch insensitive as the *eat-4(-); unc-9(-)* mutants, some non-GluCl glutamate receptors may be involved (Figure 3C).

Another line of evidence supporting the role of GluCl genes in touch response came from the finding that ivermectin treatment, which targets GluCl channels in *C. elegans*,^29^ could cause anterior insensitivity in *unc-7(-)* and *unc-9(-)* mutants but not in wild-type and *eat-4(-)* animals (Figure 3D). As an internal control, posterior touch sensitivity was not affected by ivermectin. These results confirm that the chemical synapse in the anterior touch circuit works through GluCl receptors. Lastly, to confirm that AVA is the downstream interneuron activated by the AVM-AVB pathway, we used a strain expressing the Tetanus toxin (TeTx) specifically in AVA to block its synaptic release^10^ and found that inhibiting AVA functions in *unc-9(-)* mutants but not the wild-type animals led to anterior insensitivity (Figure 3E). This result is consistent with previous findings that ablating either AVD or AVA alone did not reduce the touch sensitivity but ablating both made the animals incapable of moving backwards in response to head touch.^8^

### Developmental remodeling and lateralization of the anterior touch circuit

Given that the anterior touch circuit undergoes a developmental remodeling from the larval to adult stages after AVM joins the circuit, we monitored the potential change in the localization patterns of the electric and chemical synapses throughout development using UNC-9::GFP, INX-1::RFP, and GFP::SYD-2 expressed specifically in the TRNs. SYD-2 is a liprin-like presynaptic scaffolding protein used to label the chemical synapses.^30^ To standardize the analysis, we divided the nerve ring into three zones for anterior TRN synapses: the dorsal zone (covering only ALM synapses), the subventral zone (overlapping region for AVM and ALM), and the ventral zone (covering only AVM synapses) (Figure 4A). As the animal developed from young larvae to adults, we observed the emergence of UNC-9::GFP puncta in the ventral zone, which is expected given that the newly differentiated AVM makes gap junctions to AVD mostly in the ventral zone (Figure 4B-D).

**Figure 4.**
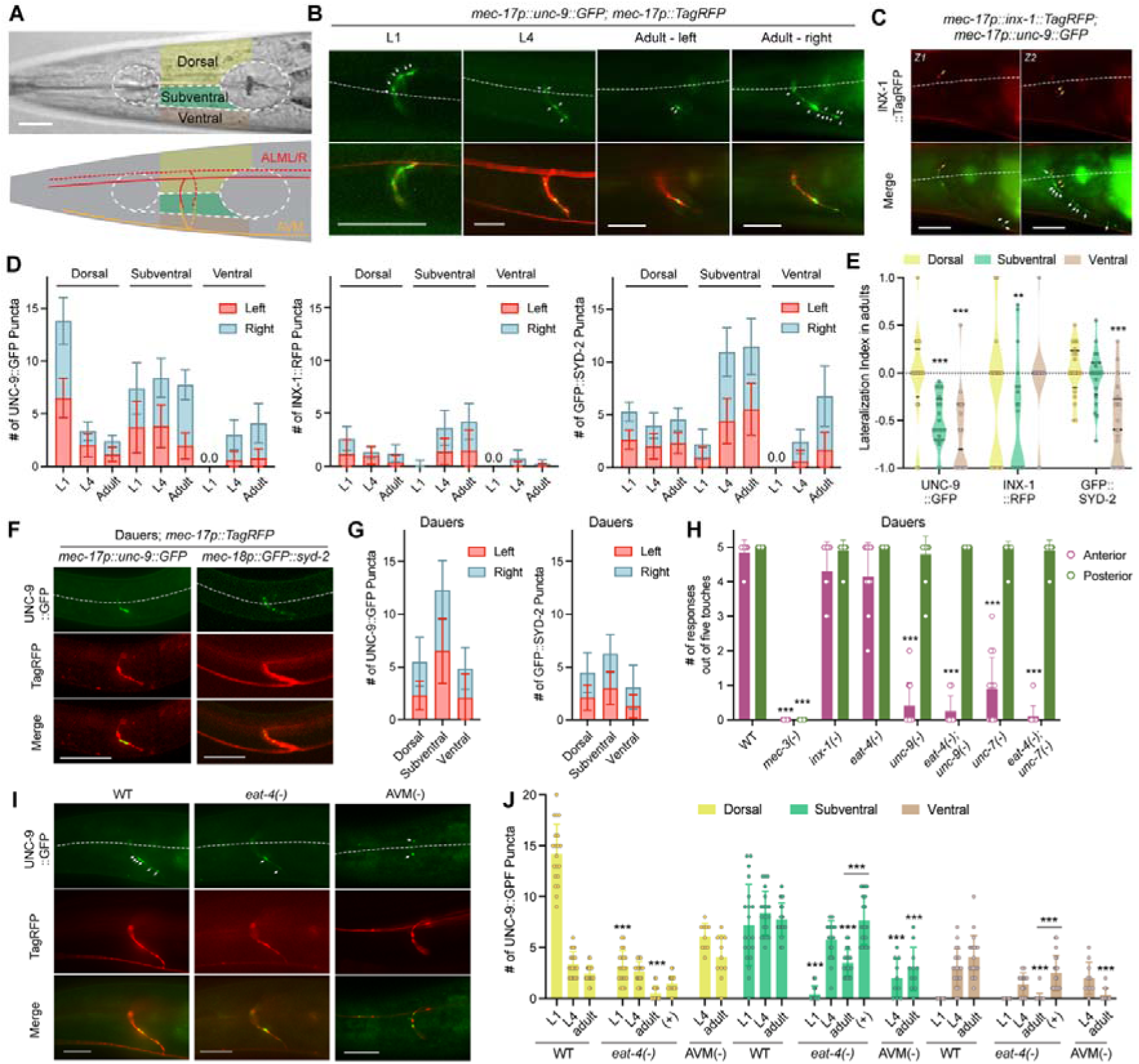
Localization of synaptic markers in the anterior touch circuit. (A) The nerve ring is divided into dorsal, subventral, and ventral zones containing ALM-only, ALM and AVM, and AVM-only synapses, respectively. (B) UNC-9::GFP signals in the TRNs at different stages. Arrows point to UNC-9::GFP puncta; white dashed lines indicate the midline separating dorsal and subventral zones. (C) In adults, INX-1::RFP colocalize with UNC-9::GFP in the dorsal and subventral zone (orange arrows), while the INX-1::RFP signal is almost absent in the ventral zone which shows abundant UNC-9::GFP puncta (white arrows). “Z1” and “Z2” indicate two focal planes of the same animal. (D) The number of UNC-9::GFP, INX-1::RFP, and GFP::SYD-2 puncta in the TRN neurites in the three zones of the nerve ring. Data are separated into left and right sides. N = 20 for each strain. (E) Violin plot of the lateralization index, defined as (# of puncta on the left side - # of puncta on the right side)/total # of puncta from both sides. Two and three asterisks indicate *p* < 0.01 and 0.001, respectively, in a one-sample *t*-test for deviation from 0. (F-G) UNC-9::GFP and GFP::SYD-2 localization in the dauer TRNs and their quantifications. (H) The number of responses out of five anterior and posterior touches in dauers of various strains. Triple asterisks indicate *p* < 0.001 in Dunnett’s test comparing mutants with the wild type. N = 20. (I-J) Images and quantifications of UNC-9::GFP signal in the anterior TRNs of *eat-4(ky5)* mutants and animals with AVM ablated. Triple asterisks in (J) indicate p < 0.001 in Tukey’s tests comparing the condition with the wild-type or between specific pairs. Mean ± SD is shown in this Figure. Scale bars = 20 μm.

Interestingly, we also observed a dramatic decrease in the number of UNC-9::GFP puncta in the dorsal zone, where ALM makes gap junctions with AVD (and also PVR) (Figure 4B-D). This developmental pruning of synapses is intriguing, since maintaining the larval ALM::AVD connection in theory can facilitate touch sensation in adults. A rather appealing hypothesis is that once the ALM::AVM and AVM::AVD gap junctions are established in late larval stages, ALM is connected with AVD through AVM, which makes the direct ALM::AVD gap junctions dispensable. Maintaining ALM::AVD gap junction may be energetically costly, given that UNC-9 was reported to turn over within 3 hours.^31^

Hints of this process were found in the early electron microscopy-based connectome studies, as it was noted that the gap junctions between AVM and AVD were easily identified in the L4 animal (JSH series) but could not be identified in sections from adults (JSE, N2U, and N2T series);^8,9,32^ the connectome of the adult male (N930 and N2Y series) did not show ALM::AVD gap junctions either.^33^ Thus, it is plausible that the ALM::AVD connection is weakened during larval development. This transition may allow AVM to consolidate the signals from ALML and ALMR and then relay the sensory inputs to AVDL and AVDR. Given that AVM can also trigger a second reversal pathway through AVB and AVA, weakening the ALM::AVD connection may enable AVM to better serve as a signaling hub for activating downstream interneurons.

We also examined the distribution of INX-1::RFP puncta, which were found mostly in the subventral zone where AVM makes connections with ALM (Figure 4C and 4D); this result is consistent with the above finding that INX-1::INX-1 gap junctions mediate ALM::AVM connection (Figure 2). Nevertheless, we also observed a few INX-1::RFP dots in the dorsal zone, which potentially label ALM gap junctions with AVD or PVR. There were generally much fewer INX-1 puncta than UNC-9 puncta, which is consistent with the genetic data that UNC-9 plays an essential role in mediating the TRN::AVD connection, while the loss of INX-1 does not affect the AVD pathway (Figure 1). Majority of the INX-1 puncta colocalized with UNC-9 puncta in the anterior TRNs (Figure S4A), likely because channels with different innexin compositions were formed at the same cell-cell junctions. For the chemical synapses, GFP::SYD-2 signals increased in the subventral and ventral zones throughout development as AVM makes various synapses to AVB, PVC, and other neurons, but the signal in the dorsal zone was stably maintained, indicating likely no pruning of chemical synapses (Figure 4D and S4B).

While examining the synaptic signals in adults, we surprisingly found left-right asymmetry in the distribution of the puncta. The right side of the AVM synaptic branch had many more UNC-9::GFP and GFP::SYD-2 puncta in the ventral zone than the left side, while there were also more UNC-9::GFP puncta on the right side in the subventral zone in adults (Figure 4D, 4E, and S4A). The few INX-1::RFP puncta also appeared to be enriched on the right side in the dorsal and subventral zones. These results suggest that both electrical and chemical synapses in the anterior touch circuit undergo a right-biased lateralization. The significance of this left-right asymmetry is unclear, although it suggests that the interneurons on the right side may receive more synaptic input than their left counterparts. In fact, AVAR is more hyperpolarized than AVAL^34^, so the activation of AVAR may require a stronger inhibition of AVBR by AVM. From the EM reconstruction data, AVM makes more chemical synapses with PVCR than PVCL (17 vs 10) and more gap junctions with ALMR than ALML (16 vs 7),^33^ which supports our findings. The same lateralization also occurs in the posterior touch circuit, where the PLMR had more UNC-9::GFP puncta than PLML (Figure S4C).

### The AVM-AVB-AVA pathway is not functional in dauers

The remodeling of the anterior circuit during larval development prompted us to analyze the touch response of dauers, which represent a developmentally arrested larval stage. The distribution patterns of UNC-9::GFP, INX-1::RFP, and GFP::SYD-2 in dauers were similar to L4, suggesting that the remodeling has occurred, as AVM has already developed its morphology and grown out the synaptic branches before entering dauers (Figure 4F and 4G; Figure S5A). However, to our surprise, the functional connectivity of the dauer circuit was similar to that of the young larvae, since the loss of the TRN::AVD gap junction in *unc-9(-)* and *unc-7(-)* mutant dauers resulted in anterior touch insensitivity (Figure 4H), like in L2 animals (Figure 1B). This result indicates that the AVM-AVB-AVA pathway is somehow not functional in dauers. However, AVM is capable of forming chemical synapses (marked by GFP::SYD-2) and delivering synaptic vesicles (labelled by GFP::RAB-3) to the synaptic branch in dauers (Figure S5B). Thus, we reason that either the dauer AVM could not release glutamate or there may be further changes in the circuit that block this pathway in dauers. Whatever the defect is, it recovers after dauer exits diapause and resumes development into adults, since the post-dauer adults of *unc-9(-)* and *unc-7(-)* mutants regained anterior touch sensitivity (Figure S5C).

### Synaptogenesis of the UNC-9 gap junction requires functional chemical synapses

Since TRNs make both electrical and chemical synapses, we examined whether the two types of synapses regulate each other. First, we found that the number of UNC-9::GFP puncta was significantly reduced in *eat-4(-)* mutants compared to the wild-type at both L1 and adult stages in all three zones (Figure 4I). The result suggests that the functional glutamatergic synapses promote the formation of gap junctions, which is consistent with similar observations made in the rat hypothalamus.^35^ Importantly, AVM-specific expression of *eat-4(+)* restored the number of UNC-9::GFP puncta in the subventral and ventral zones where AVM forms chemical synapses but not in the dorsal zone where ALM forms chemical synapse (Figure 4J). So, the regulation of gap junction synaptogenesis by chemical synapses occurs cell-autonomously. Supporting this idea, ablating AVM eliminated the ventral UNC-9::GFP signal and reduced the subventral signal but did not affect the dorsal puncta. Despite the decreased number of UNC-9::GFP gap junctions in *eat-4(-)* mutants in young larvae and adults, *eat-4(-)* mutants are touch-sensitive at all developmental stages (Figure 1B and 1C), suggesting that the remaining gap junctions are sufficient to evoke downstream motor response, and there are likely excessive UNC-9::UNC-7 gap junctions between TRNs and AVD to ensure robustness.

Conversely, we also examined the formation of chemical synapses in the absence of gap junction proteins and found that the GFP::SYD-2 puncta number and distribution were similar among wild-type, *inx-1(-)*, *unc-9(-)*, and *inx-1(-) unc-9(-)* animals (Figure S4D). It was previously reported that UNC-7 and UNC-9 regulate the differentiation of active zones at the neuromuscular junctions.^30^ We did not observe similar regulations in the TRNs.

### Stomatin-like proteins UNC-1 and UNC-24 regulate UNC-9 gap junction

We next sought to understand the regulation of the activity of the UNC-9::UNC-7 gap junction. Stomatin-like proteins were previously shown to regulate gap junctions; stomatin UNC-1 regulates UNC-9 in *C. elegans* locomotion^36^ and has recently been found to form a cap-structure that enclose the channel pore to gate the gap junction.^37^ Another stomatin-like protein UNC-24 regulates UNC-1 stability and distribution in cholesterol-rich membrane domains,^38^ where UNC-1 interacts with a sodium channel protein UNC-8.^39^ Thus, we tested the involvement of *unc-1*, *unc-24*, and *unc-8* in the touch circuit. *unc-1(-)* and *unc-24(-)* single mutants were touch insensitive in the head at L2 stage but regained the sensitivity in adults, similar to *unc-9(-)* and *unc-7(-)* mutants (Figure 5A and 5B). Just like *unc-9(-)* mutants, combining *unc-1(-)* or *unc-24(-)* mutants with either *eat-4(-)* or *inx-1(-)* mutants led to reduced anterior touch sensitivity in adults (Figure 5B), suggesting that UNC-1 and UNC-24 function in the same pathway as UNC-9. Moreover, TRN-specific expression of *unc-24(+)* in the *unc-24(-); eat-4(-)* and *unc-24(-); inx-1(-)* mutants rescued the touch response defects, confirming that UNC-24 acts in the TRNs where UNC-9 functions (Figure 5B). On the other hand, *unc-8(-)* mutants did not show touch defects in larvae.

**Figure 5.**
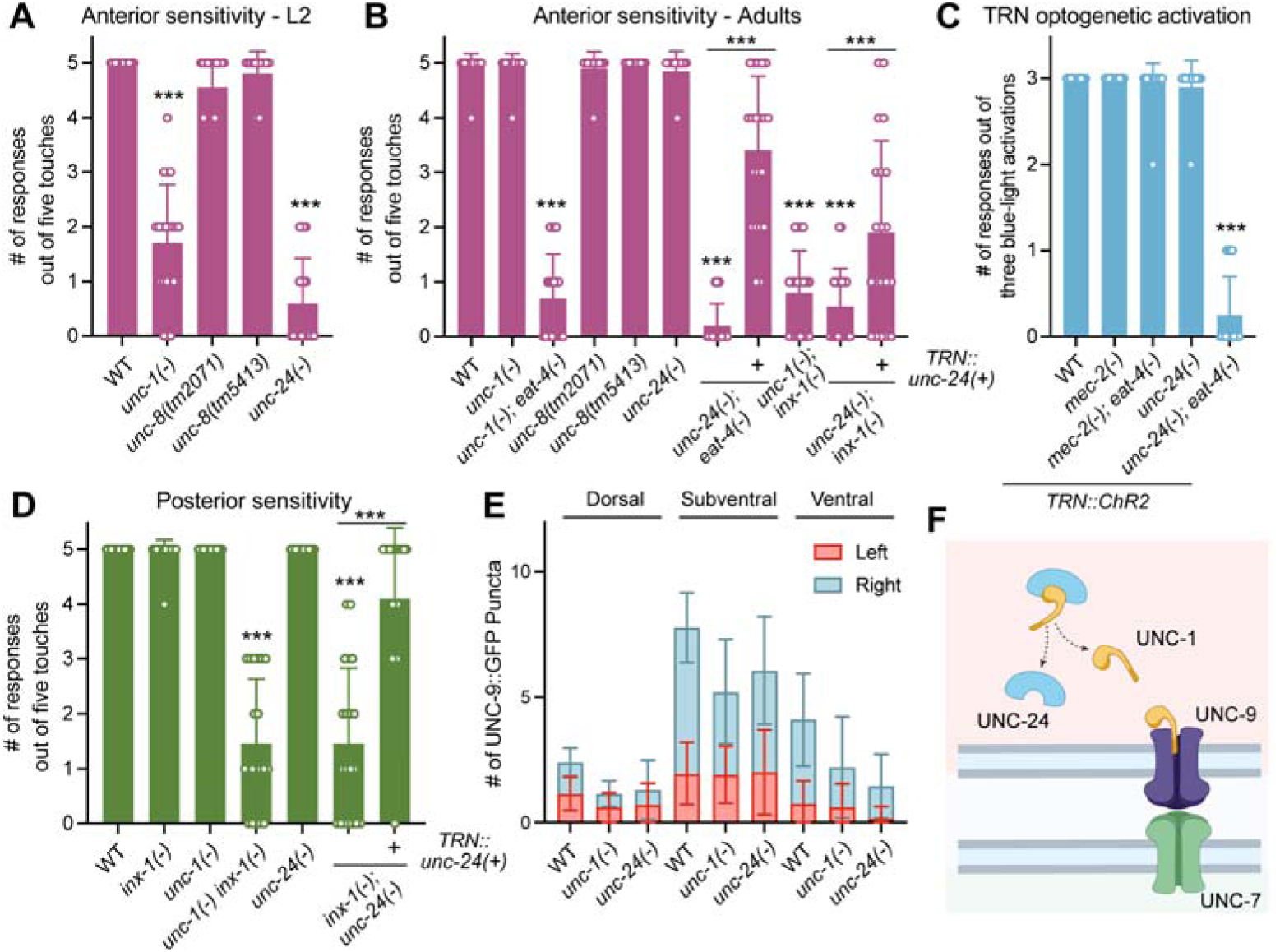
Stomatin-like protein UNC-1 and UNC-24 regulate UNC-9 gap junction. (A-B) Anterior touch responses in L2 and adult animals of the indicated strains. *unc-1(tm5728), unc-24(e138)*, *eat-4(ky5)*, *inx-1(unk29)* were used as their null alleles; *unkEx556[mec-17p::unc-24]* was used for TRN-specific rescue. Triple asterisks indicate *p* < 0.001 in Tukey’s tests comparing the mutant with the wild-type or between specific pairs. (C) The number of reversal responses out of three blue light stimulations of the head of animals carrying the *uIs94[mec-4p::ChR2::YFP]* transgene; *mec-2(u27)* allele was used. (D) Posterior touch response in adults of the indicated strains. (E) The number of UNC-9::GFP puncta in the anterior TRN neurites located in the three zones in the stomatin mutants. (F) A schematic cartoon depicting the role of UNC-24 and UNC-1 in regulating UNC-9 gap junctions. Mean ± SD is shown in this Figure. N = 20.

*C. elegans* TRNs express a third stomatin-like protein called MEC-2, which interacts with the mechanotransducer MEC-4 and is required for mechanosensation.^40,41^ To bypass the mechanotransduction step and assess the potential role of MEC-2 in regulating UNC-9, we optogenetically activated anterior TRNs and recorded the reversal response and found that both *mec-2(-)* and *mec-2(-) eat-4(-)* mutants responded to head stimulation as the wild-type animals, suggesting that MEC-2 may not regulate TRN::AVD gap junction (Figure 5C). In contrast, *unc-24(-) eat-4(-)* mutants failed to respond to the optogenetic activation of ALM and AVM, supporting a role of UNC-24 in regulating electrical synapses.

In the posterior circuit, UNC-9::UNC-7 and INX-1::INX-1 gap junctions redundantly connect PLM and PVC. We found that although *unc-1(-)* and *unc-24(-)* single mutants were touch-sensitive in the tail, *unc-1(-); inx-1(-)* and *unc-24(-); inx-1(-)* mutants show posterior insensitivity; the phenotype in the latter mutants was rescued by TRN-specific expression of *unc-24(+)* (Figure 5D). These results suggest that UNC-1 and UNC-24 only regulate the UNC-9::UNC-7 but not the INX-1::INX-1 gap junction. Furthermore, although we did not observe a significant change in the distribution pattern of UNC-9::GFP in *unc-1(-)* and *unc-24(-)* mutant adults, there appeared to be fewer puncta in the ventral zone of the nerve ring, where the TRN::AVD gap junctions were located (Figure 5E). Overall, our data indicate that the stomatin-like proteins regulate the function of specific gap junctions (Figure 5F).

### Multiple synaptic connections in the anterior touch circuit promote reversal additively

Activation of the anterior TRNs by mechanical stimuli or optogenetic excitation evokes a stereotypical reversal response which is often followed by an omega turn, and a longer reversal distance tends to have a higher chance of initiating a turn.^42–44^ So, we examined whether the various synapses in the anterior touch circuit contributes to reversal distance. First, we found that *eat-4(-)* mutants had shorter reversal distance and were less likely to initiate an omega turn after anterior touches (Figure 6A). Importantly, these defects were rescued by AVM-specific expression of *eat-4(+)*, indicating that the AVM-AVB-AVA pathway likely contributes to longer reversal distance although it is redundant in initiating the backward movement. Our finding is consistent with previous report that activated AVA release neuropeptides to increase muscles activity and promote strong reversal behavior.^45^

**Figure 6.**
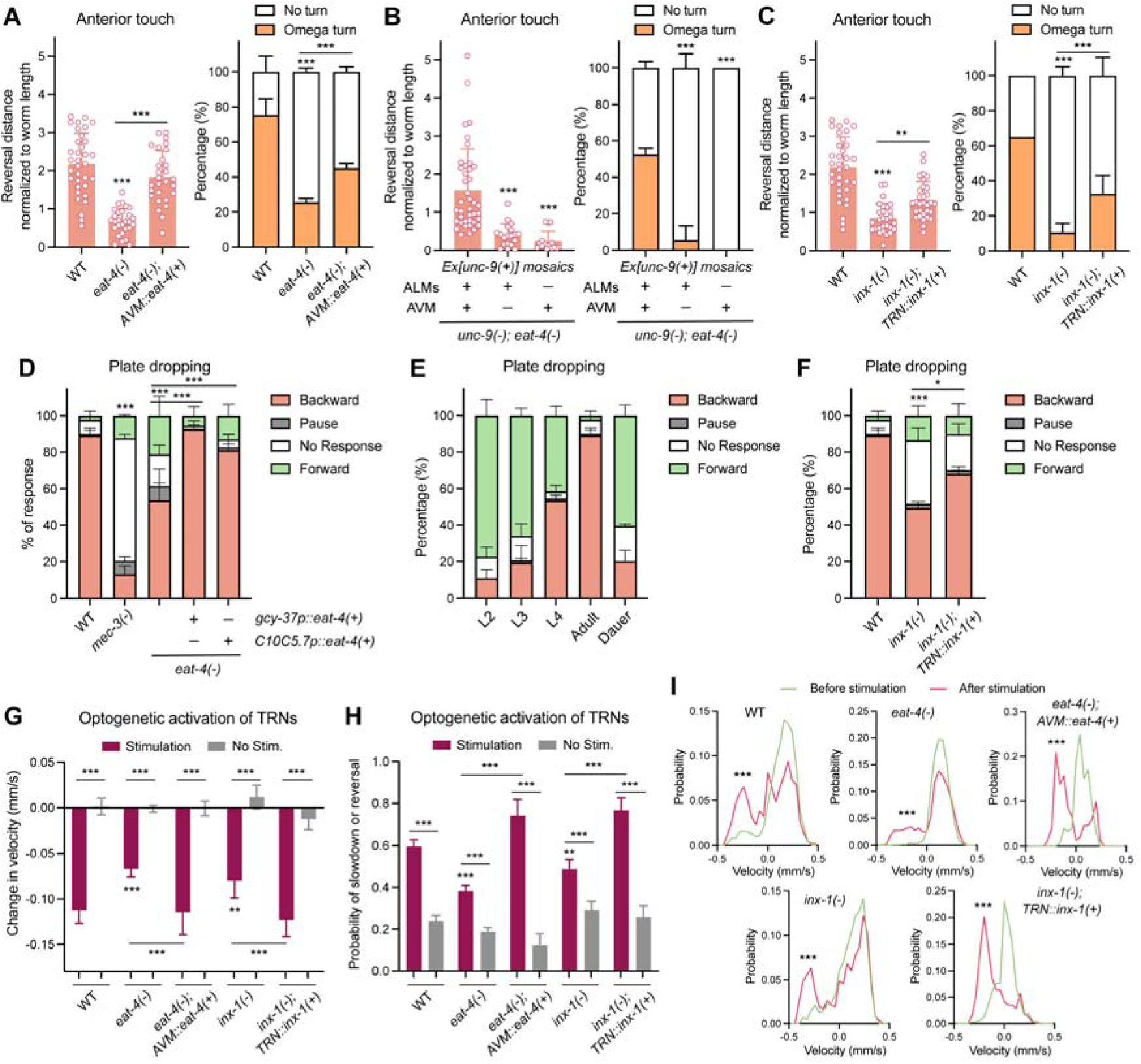
Multiple anterior synapses contribute to the touch-evoked reversal behavior. (A-C) Reversal distance (normalized to worm length) and percentage of reversals that led to an omega turn in the indicated animals triggered by anterior touch with an eyebrow. *unkEx765[gcy-37p::eat-4(+)]* was used for AVM-specific rescue of *eat-4*; *unkEx459[unc-9(+); mec-17p::TagRFP; nmr-1p::GFP]* was used for *unc-9* mosaics; *unkEx769[mec-17p::inx-1a]* was used for TRN-specific rescue of *inx-1*. Triple asterisks indicate *p* < 0.001 in Tukey’s tests comparing wild-type and mutants or selected pairs. Mean ± SD is shown; N = 30. (D) Percentage of response to plate dropping in the indicated strains; both *gcy-37* and *C10C5.7* promoters are expressed in AVM but no other TRNs. (E) Percentage of response to plate dropping of wild-type animals at different stages. (F) Percentage of response to plate dropping of *inx-1(unk29)* mutants with or without the *unkEx769* transgene. Single and triple asterisks indicate *p* < 0.05 and 0.001, respectively, in a Chi-square test. (G-H) Change in velocity and probability of slowdown/reversal upon light stimulation or no light mock stimulation for strains carrying *wtfIs458[mec-4p::Chrimson]* transgenes. CGZ2400 (N = 998 stimulation events), CGZ2399 (N = 1332), CGZ2529 (N = 153), CGZ2519 (N = 615), and CGZ2568 (N = 313) were used (see key resources table for strain details). Mean ± 95% confidence interval is shown. Double and triple asterisks indicate *p* < 0.01 and 0.001, respectively, in a Kolmogorov-Smirnov test for the left panel and a two-proportion Z-test for the right panel (both with Bonferroni correction). (I) Velocity distribution before and after the red-light stimulation. Triple asterisks indicate *p* < 0.001 in a Kolmogorov-Smirnov test. The after-stimulation velocities between *eat-4(-)* and AVM-specific rescue and between *inx-1(-)* and TRN-specific rescue show a statistically significant difference (*p* < 0.001).

Next, we found that in the *eat-4(-)* mutant adults where only TRN::AVD gap junctions mediate anterior touch response, the presence of UNC-9 in ALM and AVM both increased the reversal distance and the probability of turn, suggesting that both ALM::AVD and AVM::AVD gap junctions contribute to the reversal response (Figure 6B). INX-1-mediated ALM::AVM gap junctions also contribute to reversal distance and turn initiation although to a less extent than EAT-4 and UNC-9 (Figure 6C). These results suggest that the various synaptic connections are all required for an optimal reversal response and the subsequent omega turn that potentially changes the moving direction of the animal. For this integrated touch escape behavior, the seemingly independent AVD and AVB pathways are no longer fully redundant; instead, they work additively to promote reversal.

Another way to measure the strength of the anterior touch response is to analyze the competition of anterior and posterior responses in a plate tapping assay, in which all the TRNs are activated and the final motor output is the result of integrating anterior and posterior responses.^46^ Using a newly developed plate dropping assay in which the weight of the plate is precisely controlled (see Materials and Methods for details), we found that majority of the young adults responded by reversal as expected, whereas only ∼50% of the *eat-4(-)* mutants reversed and ∼20% even sprinted forward (Figure 6C). This result confirms that the anterior response is weakened in the *eat-4(-)* mutants. This defect can be rescued by expressing *eat-4(+)* under two promoters that are active in AVM but no other TRNs^26^ (Figure 6E), suggesting that the AVM-AVB-AVA pathway helps bias the integration of head and tail touch stimuli towards backward movement. Supporting this idea, we found that the response of the adults had much stronger bias towards reversal in plate dropping compared to young larvae that did not establish the AVM-AVB connection yet (Figure 6E). Similar to the young larvae, dauers, which have a non-functional AVM-AVB-AVA pathway, did not show a preference towards backward movement either (Figure 6E). *inx-1(-)* mutants also showed reduced probability of reversal in the plate dropping assay, and the defect could be at least partially rescued by TRN-specific expression of *inx-1(+)* (Figure 6F).

To confirm that the above effects occur at the synapse level, we used a previously established whole-field optogenetic delivery system^44^ to activate all TRNs and analyze the behavioral response via high-throughput imaging and tracking. We found that under the same light stimulation intensity, *eat-4(-)* mutants showed smaller change in velocity and a lower probability of deceleration or reversal compared to the wild-type animals, and these defects could be rescued by AVM-specific expression of *eat-4(+)* (Figure 6G-I and S6). Thus, the optogenetic data supported the above results of the touch response. We have also observed smaller change in velocity and a lower chance of slowdown and reversal in *inx-1(-)* mutants, which was rescued by TRN-specific expression of *inx-1(+)* (Figure 6G-I). This result suggests that when integrating competing anterior and posterior signals, the INX-1::INX-1 gap junctions between ALM and AVM plays a role in strengthening the anterior response and biasing the overall motor output towards reversal.

### The postembryonic AVM-AVB-AVA pathway helps adult *C. elegans* escape predation by *Pristionchus pacificus*

Finally, we explored any ecologically relevant evolutionary advantages of developing the redundant postembryonic AVM-AVB-AVA pathway. A recent study found that *C. elegans* L1 larvae are easily killed by the omnivorous nematode *P. pacificus* within a single bite, but adult *C. elegans* can survive and escapes multiple bites.^47^ We found that the TRNs appeared to play an important role in escaping from the predator, since the *mec-3(-)* and *mec-4(-)* mutants, which were touch-insensitive due to defects in TRN differentiation and loss of the mechanotransducer, respectively, were much more vulnerable to predation by *P. pacificus* than the wild-type *C. elegans* (Figure 7A). This difference was only obvious at L4 and adult stages but not in L1 and L2 animals. Interestingly, *eat-4(-)* mutants were as vulnerable as the *mec-3(-)* mutants to predation, and AVM-specific expression of *eat-4(+)* in the *eat-4(-)* mutants can at least partially increase the adult survival against *P. pacificus* predation (Figure 7B). This result suggests that the chemical synapses from AVM promote the escape of *C. elegans* from the predator likely through enhanced anterior touch response and the AVM-AVB-AVA pathway.

**Figure 7.**
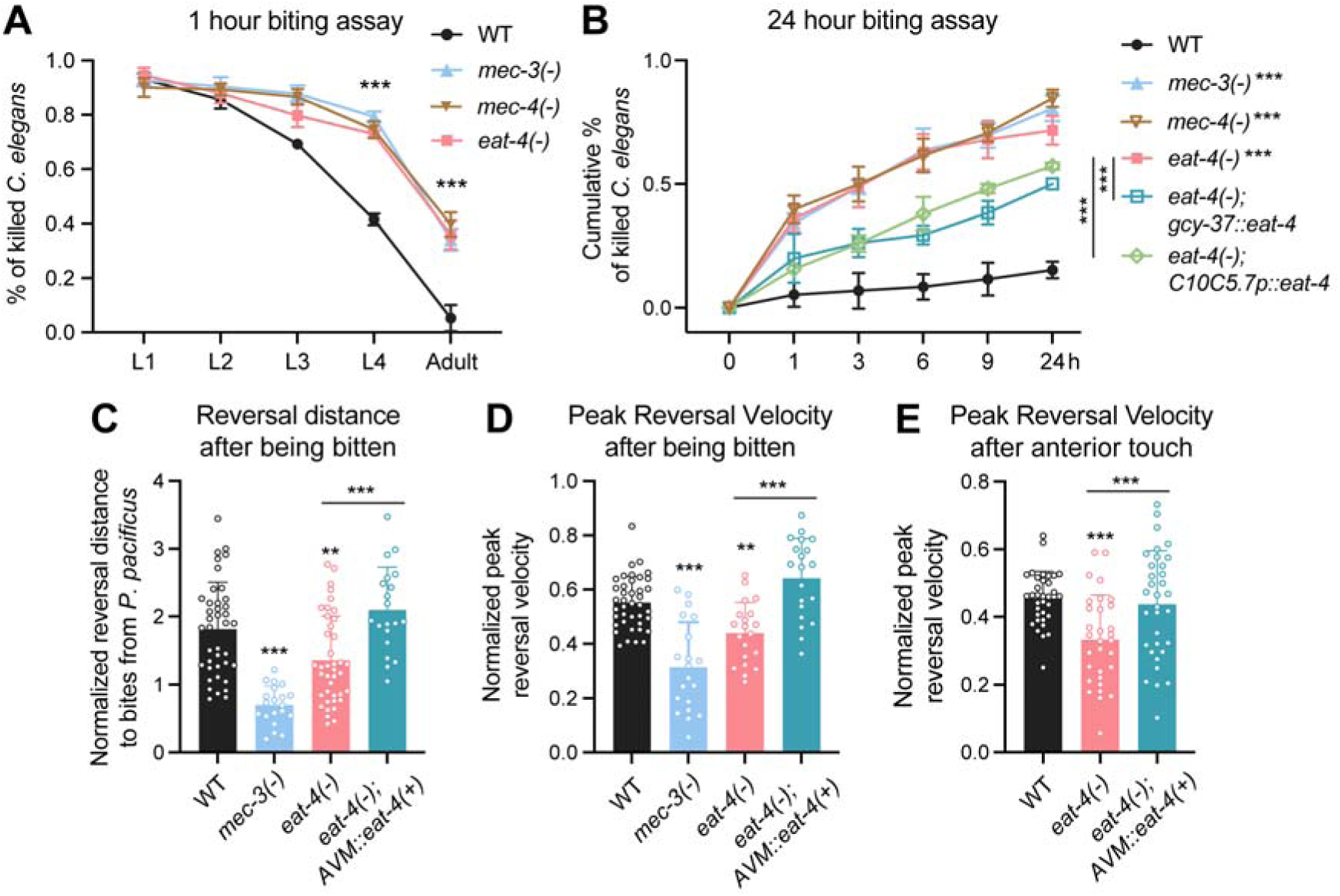
Glutamatergic synapses in the anterior touch circuit contribute to escape from predation by *P. pacificus*. (A) Percentage of *C. elegans* at various stages killed by *P. pacificus* in a one-hour biting assay. *mec-3(u184)*, *mec-4(u253)*, and *eat-4(ky5)* alleles were used. Triple asterisks indicate *p* < 0.001 in an ANOVA analysis. (B) Cumulative percentages of *C. elegans* killed by *P. pacificus* at various time points in a 24-hour biting assay. Triple asterisks indicate *p* < 0.001 in a Tukey’s test comparing the mutant and wild-type animals or selected pairs at the 24-hour time point. (C-D) Reversal distance (normalized to worm length) and peak reversal velocity of the animals after being bitten at the anterior side by *P. pacificus*. N = 40 for WT and *eat-4(ky5)*; N = 20 for *mec-3(u184)* and *eat-4(ky5); unkEx765[gcy-37p::eat-4(+)]*. Double and triple asterisks indicate *p* < 0.01 and 0.001, respectively, in a Tukey’s test. (E) Peak reversal velocity of animals after anterior touch by a hair. N = 30. Mean ± SD is shown in this Figure.

To understand why *eat-4(-)* mutants, despite being touch-sensitive, were less resistant to the predation than the wild-type *C. elegans*, we analyzed the behavioral response of *eat-4(-)* mutants upon attack by *P. pacificus* (Movies S1-S4). We found that when bitten by *P. pacificus*, *eat-4(-)* mutants escaped with a smaller reversal distance and lower reversal velocity than wild-type animals, and this difference can be removed by AVM-specific expression of *eat-4(+)* (Figure 7C and 7D). This result is consistent with the quantification of the animal’s reversal response to anterior touch by a hair (Figure 6A and 7E). Similarly, upon optogenetic activation of all TRNs, *eat-4(-)* mutants also showed smaller probability of reversal and had shorter reversal distance and duration than the wild-type animals and the phenotype was rescued by expressing *eat-4(+)* in AVM neurons (Figure S7).

Thus, the inability to reverse fast enough and far enough (to trigger an omega turn that changes the direction of the animal) may render the *eat-4(-)* mutants easier targets than the wild-type animals when encountering *P. pacificus*. The AVM-AVB-AVA pathway, seemingly redundant in mediating a simple touch avoidance, plays a crucial role in promoting and sustaining the touch-evoked reversal and therefore facilitating survival in complex environments involving predators.

## DISCUSSION

As one of the first connectomes fully mapped at the single cell resolution,^8^ the touch response circuit represents a classical example of sensory-motor reflex. By characterizing the molecular basis of the neuronal connection in the circuit, we uncovered a common theme of redundancy for the connection between sensory neurons and command interneurons, which enables robust response of the circuit. Strikingly, the redundancy can happen at multiple levels. First, the anterior circuit in adult animals possesses two redundant neural pathways (TRN::AVD gap junctions and AVM-AVB-AVA chemical synapses) that led to the activation of two different interneurons that control backward movement. Second, in the posterior circuit, the two distinct gap junction channels (UNC-9::UNC-7 and INX-1::INX-1) redundantly regulate the connection between posterior sensor PLM and the interneuron PVC that controls forward movement. Third, at the AVM to AVB chemical synapses, the postsynaptic receptors (i.e., the glutamate-gated chloride channels) show a high level of redundancy at the gene level. Thus, diverse strategies were used to build redundancy at the neural pathway, synaptic, and molecular levels. Such versatility may reflect the need to adapt to varying environmental stimuli during the evolution of the touch reflex in *C. elegans*.

Our work also reveals insights about the use of invertebrate innexin in constructing gap junctions. First, our genetic evidence suggests that the synaptic partners can use different innexins to form independent gap junction channels. For example, PLM uses UNC-9 and INX-1 hemichannels to form gap junction channels with UNC-7 and INX-1 hemichannels on PVC, respectively. The UNC-9::UNC-7 channel does not require INX-1 and vice versa, although their localizations were entirely overlapping under the light microscope, suggesting no clustering of distinct channels. This finding suggests a pairing rule that allows specific innexins to recognize each other when constructing electrical synapses. We suspect similar principles also exist for the vertebrate connexins. Second, there can be one dominant and functionally critical hemichannel pairing at the cell-cell junction. Both TRNs and AVD express *inx-1*, but INX-1 does not seem to be functionally important as UNC-9 and UNC-7 in mediating the TRN::AVD connection. For example, unlike *unc-9(-)* and *unc-7(-)* mutants, *inx-1(-)* mutants are not touch-insensitive in the larvae. We reason that this functional difference may be due to high expression or protein stability (because of auxiliary factors) of UNC-9 and UNC-7 compared to INX-1, which results in more abundant UNC-9::UNC-7 channels than INX-1::INX-1 channels at this connection. Third, heterotypic gap junction channels may have directionality, as UNC-9::UNC-7 channels may only allow signal transduction from the UNC-9 to the UNC-7 side.

Previous studies suggest that the hemichannel UNC-7 may exist in TRNs to serve as a mechanosensitive channel, and *unc-7* knockdown led to reduced calcium signal in response to mechanical stimulation.^48^ In our hands, *unc-7(-)* mutant adults are completely touch-sensitive, suggesting that even if there is a role of UNC-7 in mechanosensation, it is likely to be very minor. Conflicting results about the expression pattern of *unc-7* were also shown in the literature, as fosmid reporters showed no *unc-7* expression in the TRNs of well-fed hermaphrodites.^16^ Our mosaic analysis supports that UNC-7 functions in the AVD interneurons to promote touch response and is not required in the TRNs.

Our work also highlights a critical role of AVM as a signaling hub in the anterior circuit. As a late comer, the joining of the postembryonic AVM into the circuit leads to significant remodeling of the circuit. First, the original ALM::AVD gap junctions appear to be weakened when AVM forms connections with AVD; second, AVM builds a second neural pathway for backward movement by making chemical synapses with the AVB interneuron; third, AVM connects both ALML and ALMR through gap junctions to solidate lateral signals. Fourth, the AVM connections also appear to be lateralized with more synapses on the right synaptic branch than the left one. The significance of such developmental remodeling is not entirely clear. One purpose may be to strengthen the anterior response, so when integrating opposing signals from the head and tail, the overall motor output favors backward movement in late larvae and adults. The strengthened anterior response may also help escape predators at the adult stage.

Interestingly, the genes that were seemingly redundant in a simple touch response assay showed their indispensability in more sensitive assays that measure the extent of the response (e.g., reversal distance, velocity, and chance of initiating a turn). The results of these assays suggest that the ALM::AVM, ALM::AVD, and AVM::AVD gap junctions and the AVM-AVB chemical synapses promote reversal response in an additive manner, which sustains the reversal long enough so that the animals can enter an omega turn to change the direction of movement. We suspect that this change of direction may be important for escaping predation.

Studies from diverse organisms indicate that redundancy is likely a common strategy to ensure robustness in sensory-motor reflexes. This redundancy can happen at the molecular sensor level (e.g., three TRP channels redundantly mediate the sensing of acute noxious heat in mice)^49^, at the sensory cell level (e.g., the retinal ganglion cells are redundant in coding visual information in salamander)^50^, at sensory organ level (e.g., superior tentacles and inferior tentacles are redundant in order aversion in slugs)^51^, at the signal-relaying neuron level (e.g., multiple medulla projection neurons redundantly mediate color vision in Drosophila)^52^, and at the neural network levels (e.g., redundant circuits in the commissural pathway in primates).^53^ Therefore, there is likely an evolutionary advantage in building redundancy in the basic circuit unit of the nervous system.

### Limitations of the study

Although we used calcium imaging to analyze neuronal activities in the circuit of immobilized animals, we could not visualize neural signal propagation in the circuit of free-moving animals upon touch by a hair or optogenetic activation of TRNs. Being able to monitor the calcium signal in both TRNs and interneurons in various mutants of synaptic proteins can help further elucidate the role of each neuronal connection in the circuit. Moreover, we do not understand the mechanisms governing the developmental pruning of the dorsal synapses in the anterior circuit, the lateralization of the AVM synapses, and the regulation of gap junction by EAT-4. Further studies are needed to explore these mechanisms. We also failed to identify the non-GluCl glutamate receptors that mediate the AVM-AVB synapses. Finally, it is unknown whether the postembryonic AVM-AVB pathway also contributes to other predation events, such as the predation by predacious fungi, although *mec-4(-)* mutants appear to be more vulnerable to the trapping by these fungi.^54^

## RESOURCE AVAILABILITY

### Lead contact

Further information and requests for resources and reagents should be directed to and will be fulfilled by the lead contact, Chaogu Zheng.

### Materials availability

Strains and plasmids generated in this study will be shared by the lead contact upon request.

### Data and code availability

1. The raw behavioral data have been deposited to Figshare and are publicly available as of the date of publication. Accession numbers are listed in the key resources table.
2. The code used to collect and analyze the behavioral dataset is available at Github and is publicly available. Links are listed in the key resource table.

## ACKNOWLEDGEMENTS

We thank Kelly Wing Ka Lo in the Zheng lab for technical assistance and Shuting Han in the Chalfie lab for helping with the *eat-4(-)* genetic screen. We thank Prof. Ralf Sommer for providing the *P. pacificus* strain RS5194. This work is supported by the National Natural Science Foundation of China (Excellent Young Scientists Fund for Hong Kong and Macau, 32122002), and the Research Grant Council of Hong Kong (GRF 17105523, GRF 17106322, GRF 17113324, and CRF C7026-20G). A.M.L. is supported by the National Science Foundation through an NSF CAREER Award (IOS-1845137) and the Simons Foundation award SCGB 543003. This work also received support from National Institutes of Health grants GM30997 and GM122522 to M.C. Some strains were provided by the Caenorhabditis Genetics Center, which is funded by the NIH Office of Research Infrastructure Programs (P40 OD010440), and the National BioResource Project (NBRP), which is funded by the Japanese government. Some figure panels were created using Biorender.com.

## AUTHOR CONTRIBUTIONS

C.Z., M.C., and H.H. conceived the study. H.H. and E.K.H.F. carried out most of the experiments and data analyses. S.K. conducted the optogenetic stimulation experiments. H.M.T.L. helped construct transgenic animals. A.M.L. supervised the optogenetic studies. C.Z. and H.H. prepared the manuscript. All authors read and approved the manuscript.

## SUPPLEMENTAL INFORMATION

**Document S1. Table S1-S3, Figure S1-S7**

**Movie S1. An example video of *P. pacificus* biting wild-type *C. elegans* and its subsequent escape response.** Adult animals of *P. pacificus* RS5194 and *C. elegans* N2 strains were used. The animals were labelled by white circles in the first frame.

**Movie S2. An example video of *P. pacificus* biting *mec-3(-) C. elegans* and its subsequent escape response.** Adult animals of *P. pacificus* RS5194 and *C. elegans* TU656 *mec-3(u184)* strains were used. The animals were labelled by white circles in the first frame.

**Movie S3. An example video of *P. pacificus* biting *eat-4(-) C. elegans* and its subsequent escape response.** Adult animals of *P. pacificus* RS5194 and *C. elegans* CGZ202 *eat-4(ky5)* strains were used. The animals were labelled by white circles in the first frame.

**Movie S4. An example video of P. pacificus biting eat-4(-); Ex[AVM::eat-4(+)] C. elegans and its subsequent escape response.** Adult animals of *P. pacificus* RS5194 and *C. elegans* CGZ2370 *eat-4(ky5) III; unkEx765[gcy-37p::eat-4(+); mec-17p::TagRFP; myo-2p::mCherry]* strains were used. The animals were labelled by white circles in the first frame.

## STAR □ METHODS

### KEY RESOURCES TABLE

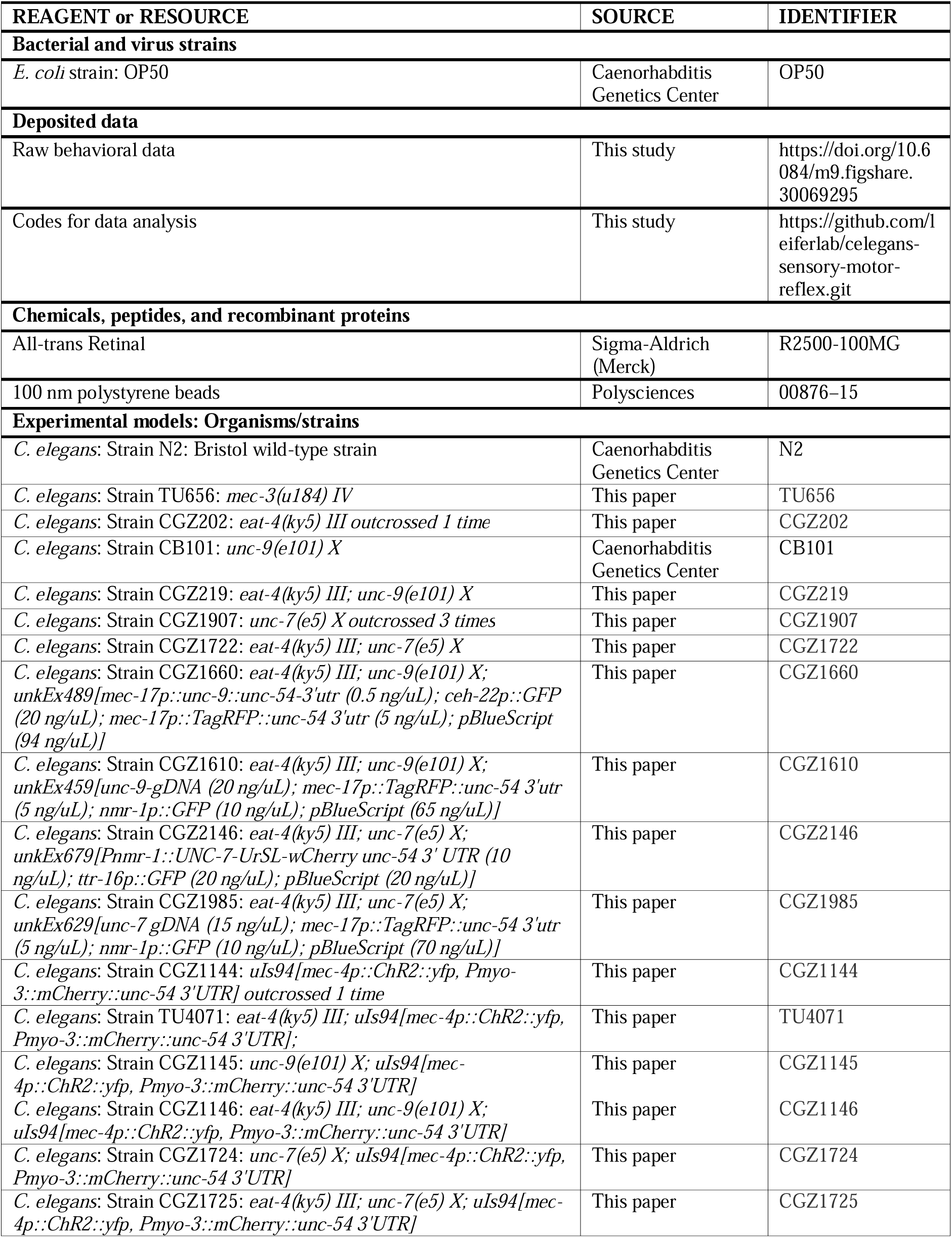
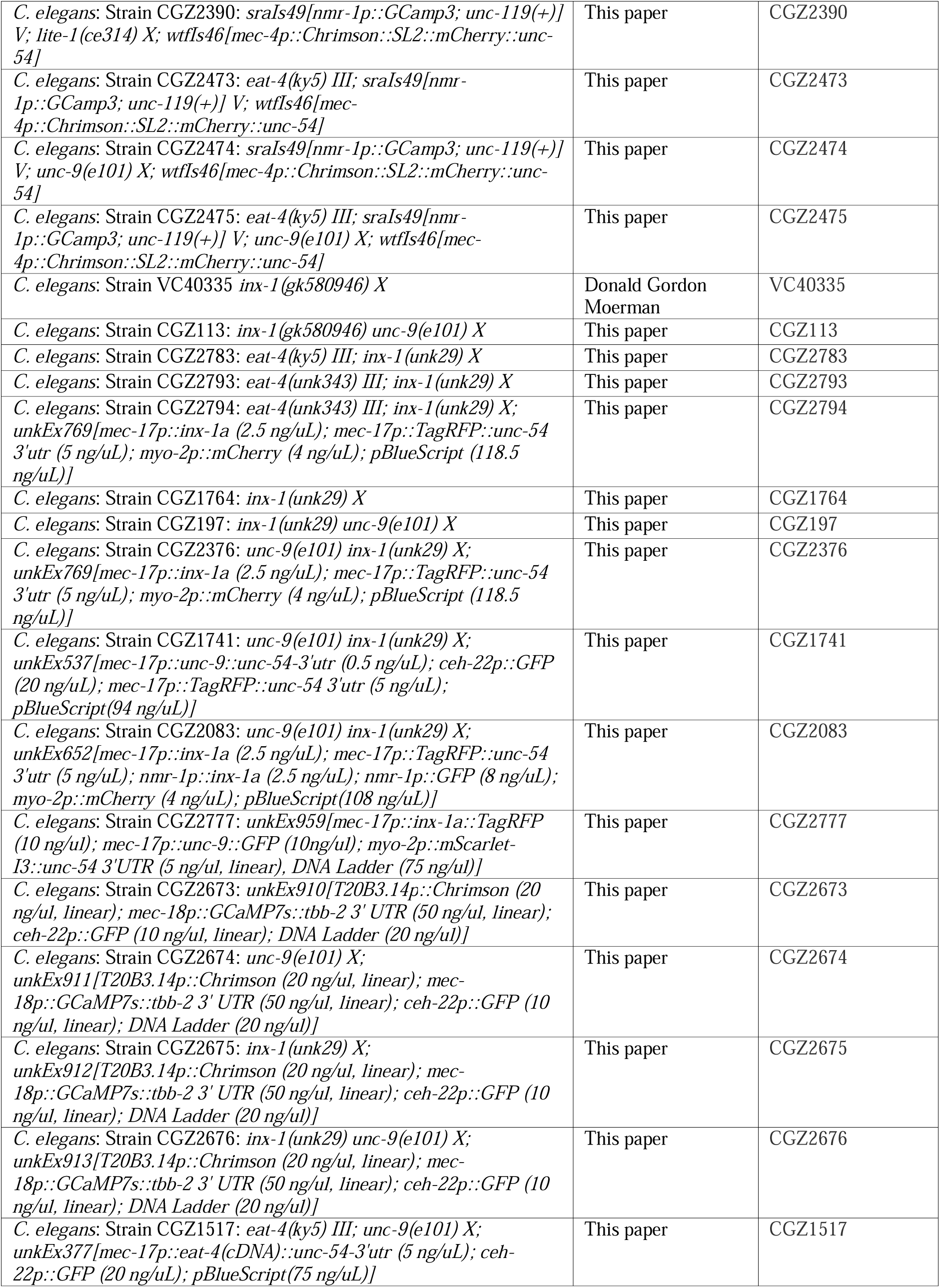
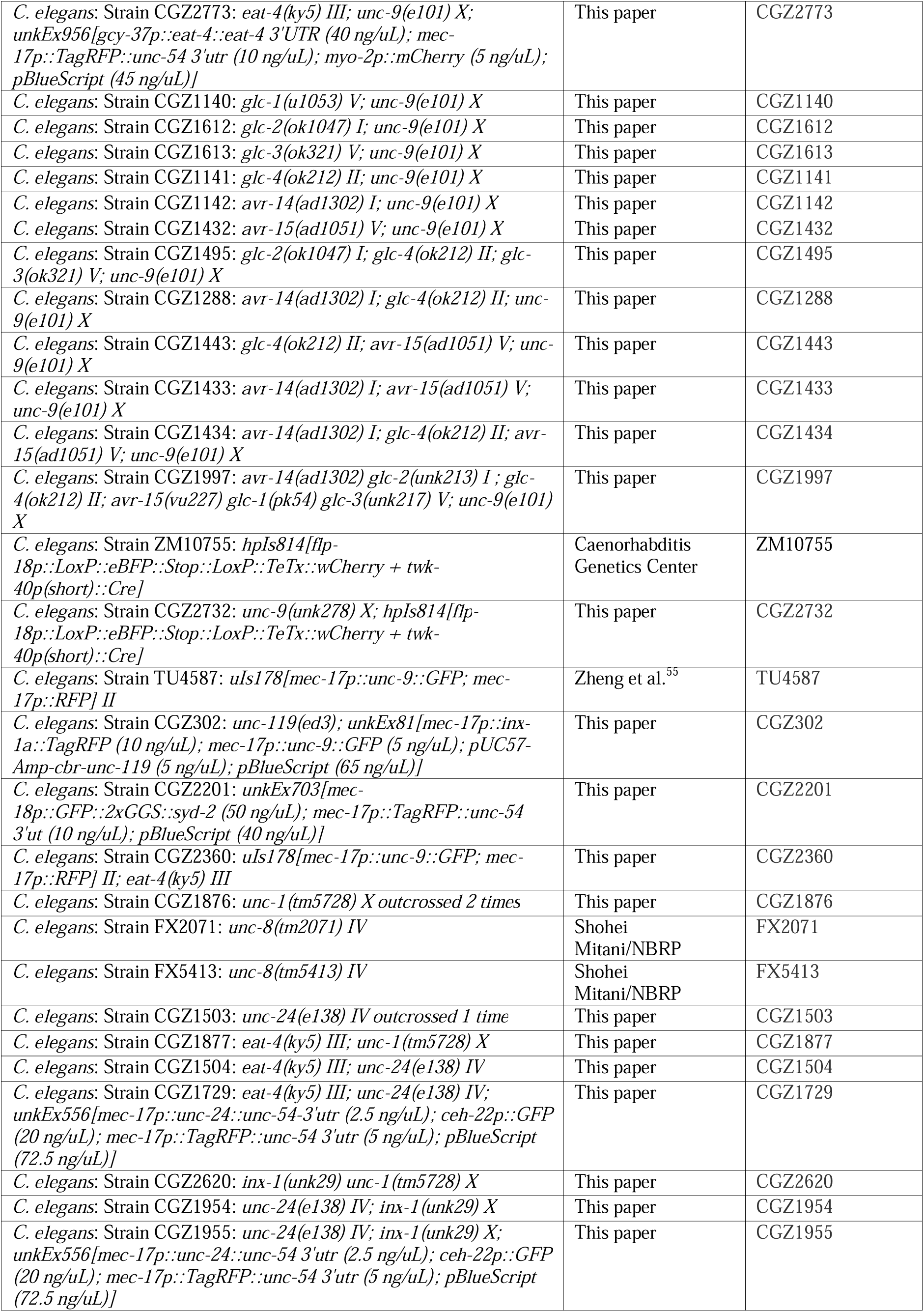
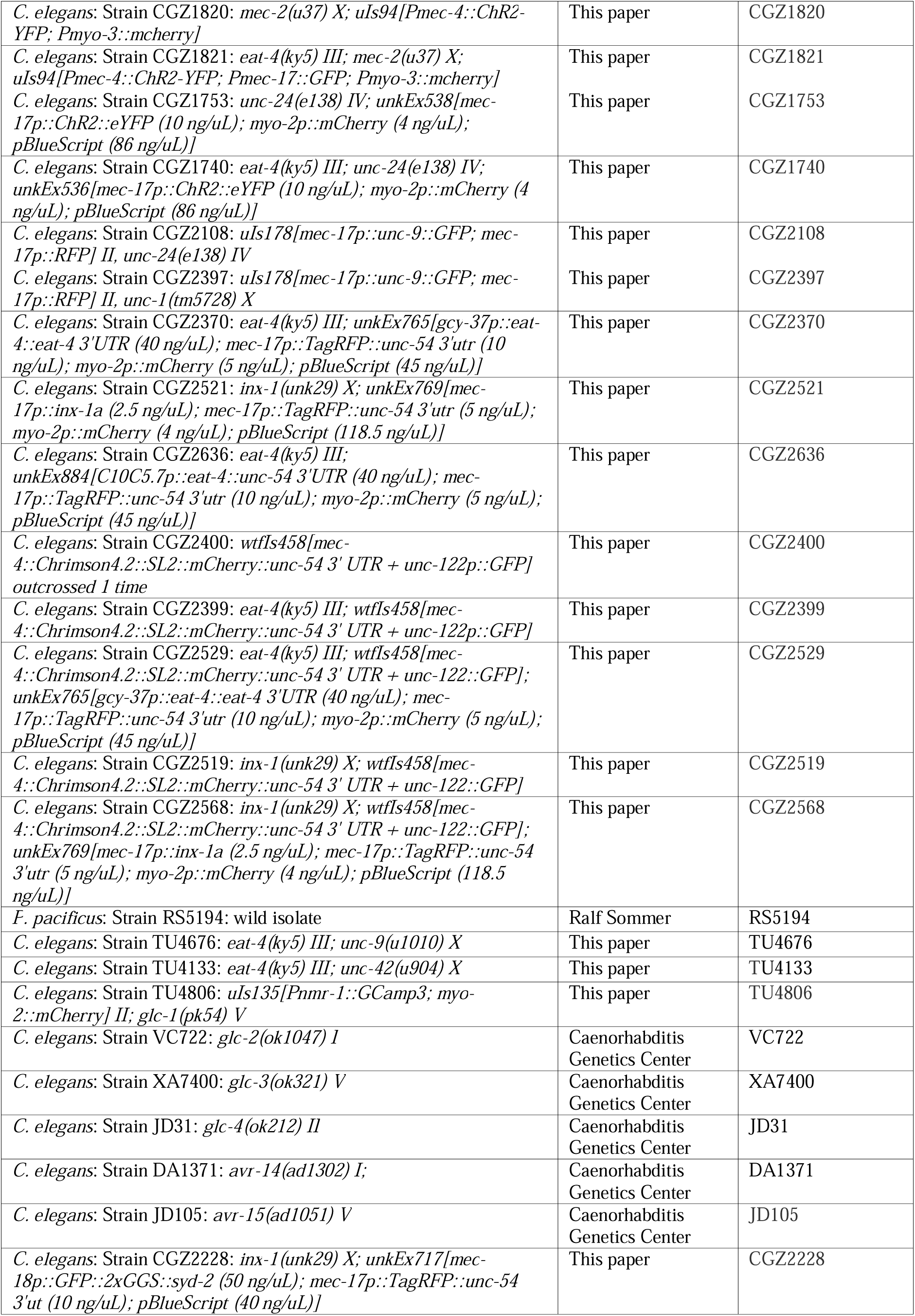
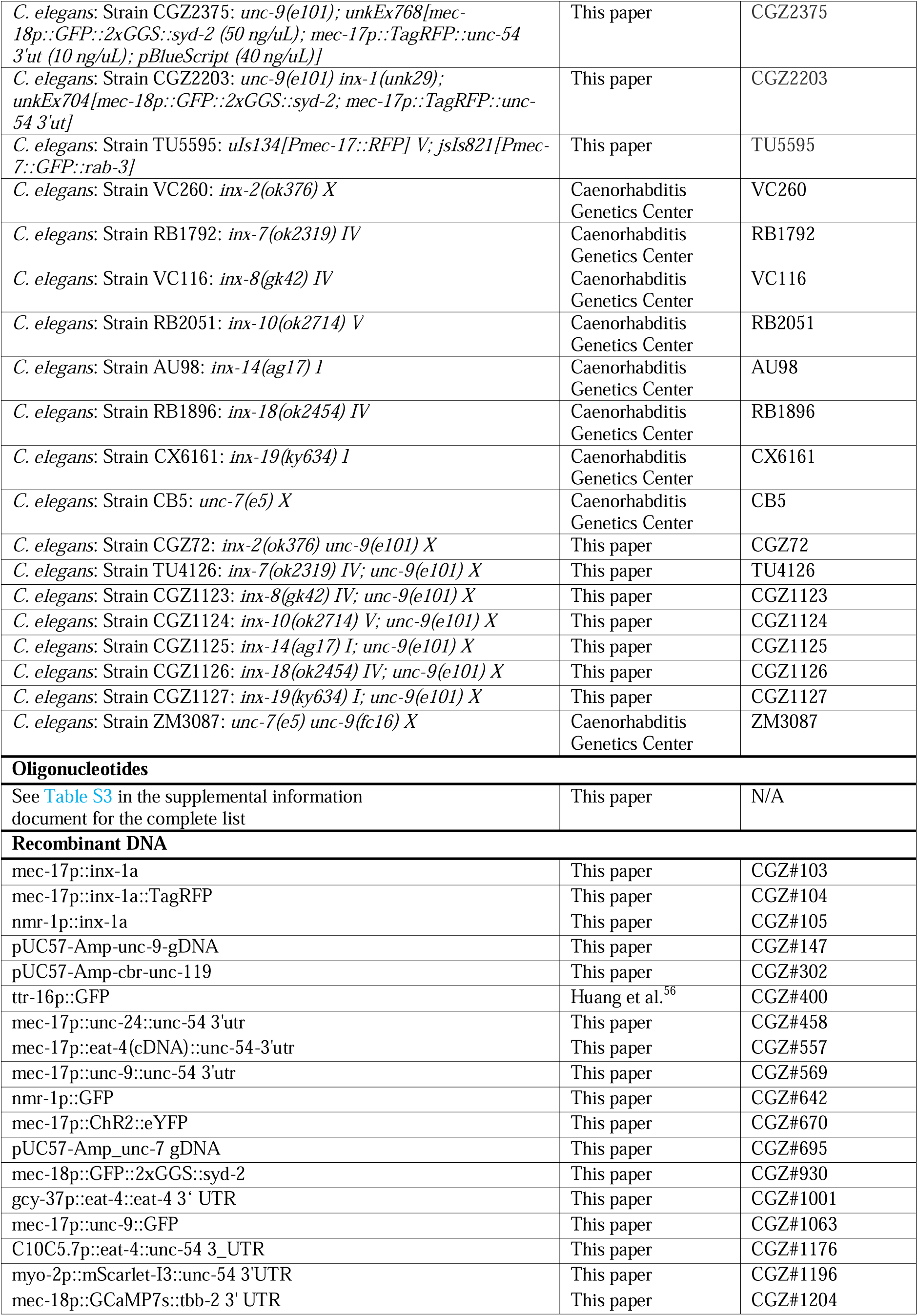
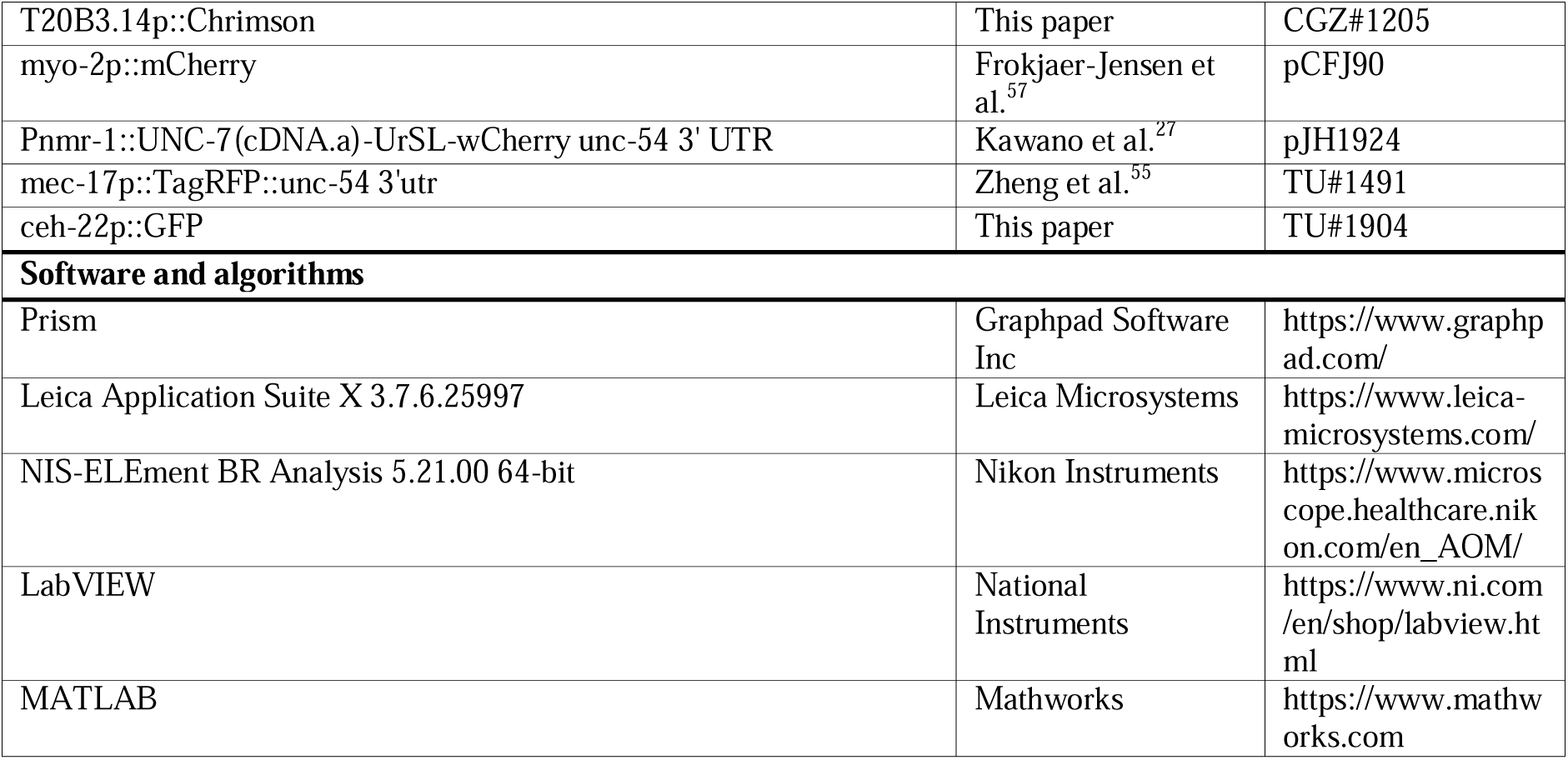

## EXPERIMENTAL MODEL AND STUDY PARTICIPANT DETAILS

### C. elegans strains

*C. elegans* wild-type (N2), mutant, and transgenic strains were maintained as previously described.^58^ Most of the experiments were performed at 20°C on NGM plates seeded with *E. coli* (OP50) as a food source unless otherwise indicated. *P. pacificus* wild-type RS5194 strain was a generous gift from Prof. Ralf Sommer at Max Planck Institute for Biology Tübingen and was cultured according to standard protocols.^59^ Some strains were obtained from either the Caenorhabditis Genetics Center or the Japanese National BioResource Project, and others were created in this study. All strains used in the experiments are listed in the key resources table.

## METHOD DETAILS

### DNA constructs and transgenesis

Most of the plasmids used in this study were constructed using a pUC57 backbone. To achieve TRN-specific gene expression, *inx-1a*, *unc-9*, and *unc-24* coding sequences were cloned from genomic DNA and then inserted between a 1.9-kb *mec-17* promoter and *unc-54* 3’UTR using Gibson Assembly (ClonExpress, Vazyme, China). The resulted constructs were injected into the corresponding *inx-1(-)*, *unc-9(-)*, and *unc-24(-)* mutants to form transgenic animals with extrachromosomal arrays. To perform interneuron-specific rescue, we cloned *inx-1a* from genomic DNA and ligated it between a 5.1 kb *nmr-1* promoter and upstream of the *unc-54* 3’UTR.

For mosaic analysis, an 8.9-kb *unc-9* coding fragment containing its native promoter and UTRs was cloned into a pUC57 vector using the primers 5’-cgtcgtactttatgtgctcgtgc-3’ and 5’-caggaaacagctatgaccatatggcaaagtttgtgggttcc-3’. The plasmid was then co-injected with TRN markers (*mec-17p::TagRFP*) and interneuron markers (*nmr-1p::GFP*) into *unc-9(e101)* mutants to form an extrachromosomal array. Animals with the array present or absent in TRNs or interneurons were picked and used for various assays. Similarly, a 10.2-kb *unc-7* coding fragment with its native promoter and UTRs was cloned into a pUC57 using the primers 5’-gtgaacttatgatcctcaaatagaattttttaggacctc-3’ and 5’-gttgtaaaacgacggccagtcggcattccagaaccgaaac-3’ and co-injected with TRN and interneuron markers into *unc-7(e5)* mutants.

To label chemical synapses, GFP was linked to the N terminus of SYD-2 with a 2xGGS linker^59^ and expressed under a 0.4-kb *mec-18* promoter. To label INX-1, its coding sequence was ligated downstream of a 1.9-kb *mec-17* promoter and upstream of the TagRFP coding sequence and *unc-54* 3’UTR. The resulted constructs were injected into worms to form extrachromosomal arrays. *uIs178[mec-17p::unc-9::GFP; mec-17p::RFP]*^55^ was used to visualize UNC-9 localization.

Chrimson coding sequence was cloned from *mec-4p::Chrimson::SL2::mCherry::unc-54* (Addgene, Plasmid #107745) and ligated downstream of a 3.8-kb *T20B3.14* promoter for ALM::Chrimson expression. *C. elegans*-codon optimized GCaMP7s coding sequence was synthesized by Tsingke Biotechnology (China) and ligated downstream of a 0.4-kb *mec-18* promoter. The resulted constructs were injected together to create the *unkEx911[T20B3.14p::Chrimson; mec-18p::GCaMP7s; ceh-22p::GFP]* transgene for Calcium imaging studies. All transgenes, alleles, and strains used in this study are listed in the key resources table.

### CRISPR/Cas9-mediated gene editing

To delete *inx-1*, single-guide RNA (sgRNAs) targeting 5’-CCGGAATGCTTCTATATTAT-3’ in exon 1 and 5’-TGTGGAGAGGACTGCTGCAC-3’ in exon 3 were generated using the NEB EnGen sgRNA Synthesis Kit (E3322V) and injected together with recombinant Cas9 (EnGen S. pyogenes Cas9 NLS from NEB, M0646T) into C. elegans gonads. The consequent transformants were screened for any deletion in *inx-1*. Allele *unk29* was isolated as a presumably null allele with a 490-bp deletion covering its start codon and exons 1 to 3.

Similarly, *glc-1* was deleted using sgRNAs targeting 5’-gcaATGGCTACCTGGATTGT-3’ in exon 1 and 5’-GGACAGTTCCACTGAATCGC-3’ in exon 5. The resulted *u1053* allele is a 1952-bp deletion removing most of the coding sequences. *glc-2* was knocked out by sgRNAs targeting 5’-cctcctcttcctccatatcacat-3’ in its promoter and 5’-CCTCCGGCATTTCTCACAGTCCC-3’ in exon 2. The presumably null allele *unk213* with a 670-bp deletion covering its start codon and exons 1 to 2 was isolated. *glc-3* was knocked out by sgRNAs targeting 5’-ccgtctctggaggtcacatc-3’ in its promoter and 5’-ctgttccgcaatcataacca-3’ after exon 1. A 232-bp fragment covering its start codon and exons 1 was deleted and replaced by an extra T in allele *unk217*.

*unc-9* was knocked out by sgRNAs targeting 5’-CTGTACACCGTGAACATCGT-3’ in exon 7 and 5’-GAATCGAGGGATATAAAATGG-3’ in exon 8. Allele *unk278* with a 504-bp deletion covering exons 7 to 8 and a 20-bp addition was isolated. *eat-4* was knocked out by sgRNAs targeting 5’-CACAGCAAATgtaagtgtat-3’ in exon 2 and 5’-AATAGATGAGCTTAGTGTCA-3’ in exon 4 as previously described.^26^ Allele *unk343* with a 455-bp deletion covering exons 2 to 4 was isolated.

### Forward genetic screen

The *eat-4(ky5)* mutants were used as the starter strain for the forward genetic screen. An eyebrow hair was used to touch the F2 progeny of mutagenized P0 animals, and mutants with defective anterior touch response but normal posterior touch sensitivity were isolated. We screened ∼5,000 haploid genomes and identified two such mutants; both showed Unc phenotypes. Targeted Sanger sequencing of the *unc-7* and *unc-9* loci revealed that one mutant (*u1010*) carries a nonsense mutation (Q334*) in *unc-9*. Complementation tests showed that the other mutant (*u904*) failed to complement *unc-24(e270)*; so, we reason that *u904* is an *unc-24* mutant allele.

### Gentle touch assay

Gentle touch experiments were conducted on synchronized day-one adult hermaphrodites following the standard protocol.^8,60^ The testers were blind to the sample identities during the experiments. An eyebrow hair was used to stroke across the body just behind the pharynx for anterior touch response or just before the anus for the posterior touch response. The anterior or the posterior body of each worm was touched five times, and a response was counted when the worm moved away from the eyebrow after the touch. In most of the experiments, 20 worms were used. Less worms (usually >10) were used for mosaic analysis due to the difficulties of picking the animals with desired mosaic patterns. Larvae at the L2 and L3 stages are much smaller than adults, so the standard gentle touch protocol for adults may generate too strong stimuli for the larvae. We adopted a gentler stimulation protocol by only touching the side of the animals with the eyebrow hair. This method was previously used to study subtle differences in touch sensitivity.^61^

For mosaic analysis, animals carrying the extrachromosomal array were observed under a fluorescent stereomicroscope for the presence of fluorescent markers in TRNs or interneurons. Animals with desired mosaic patterns were picked and transferred to a new plate and rested for at least one hour before the touch test.

### Quantification of reversal response

Behaviors of worms were recorded by either a Nikon SMZ745T stereomicroscope with a CTS CWHC-8MB 4K microscope camera or an Olympus SZX16 stereomicroscope with a Nikon FI3 camera. Videos were analyzed using the software NIS-ELEment BR Analysis 5.21.00 64-bit.

Reversal distance was defined by the distance covered from the time the animal started moving backward to the time the animal stopped moving backward. The distance was then normalized by the length of the worm. An omega turn was defined as any turn that causes the direction to deviate more than 90 degrees from the initial bearing before the reversal. Turns greater than 90 degrees where the worm’s head did not touch the body were considered open omega turns. Turns where the head touched the body were considered closed omega turns.^45^ We included both open and closed turns as omega turns. The peak reversal velocity during the reversal phase was defined as the normalized reversal distance within the first second of a stimulus.^45^

### Dauer Experiments

Dauers were generated by growing the worms under high-density and food-scarce conditions at 25°C and then selecting the dauers through a 1% SDS treatment according to previous methods.^16,62^ In detail, worms were first bleached, and the resulted eggs were placed on NGM plates seeded with OP50 at 20°C. Two days later, ten L4 animals were picked and transferred to 60-mm NGM plates with OP50, and the plates were stored at 25°C. Dauer worms usually appeared after one week. On the day of the experiment, worms were washed off the plates with M9 and collected in a 1.5-mL centrifuge tube by centrifugation. 1% SDS was added to wash the worms once. Then, worms were incubated in 1% SDS solution for 30 minutes with shaking, and only dauers could survive the incubation. After the treatment, worms were washed twice with M9 and placed on empty NGM plates without food and were allowed to recover at 20°C for at least 3 hours before the touch tests or imaging experiments.

### Optogenetics with Channelrhodopsin

TRN optogenetic activation-induced avoidance assays were performed based on the classic protocol.^63^ Worms expressing Channelrhodopsin-2 in TRNs from the *uIs94[mec-4p::ChR2::yfp, Pmyo-3::mCherry]* transgene^61^ were bleached, and the resulted eggs were placed on a bacterial lawn containing 500 μM all-trans retinal (R2500-500MG; Sigma).

Plates with all-trans retinal were wrapped in aluminum foil and stored at 20°C. Three days later, worms that reached the day-one adult stage were illuminated with a momentary blue light (450-490 nm) at their heads using a custom-made fluorescent stereomicroscope. Each worm was activated three times, and a response was counted only when a worm moved backward within 0.5 second of shining the light on its head. 20 animals were tested for each strain.

### Optogenetics with Chrimson and high-throughput behavioral analysis

A previously established method was used to stimulate the animals carrying the *wtfIs458[mec-4::Chrimson4.2::SL2::mCherry; unc-122::GFP]* transgene through whole-field illumination.^26,64^ Briefly, bleached eggs were hatched in M9 overnight and the synchronized L1 larvae were transferred to freshly seeded plates containing 1 ml of 0.5 mM *all-trans*-retinal (Cat. # R2500, Sigma-Aldrich) mixed with OP50 and stored in the dark at 20°C until day-one adult stage. On the day of the experiment, the young adult worms were washed in M9 and transferred to an empty agarose plate for experiments. Excess M9 solution was absorbed with a Kim wipe before collecting the behavioral data as described previously.^65^

An agarose plate (without food) containing 40-50 freely moving worms was illuminated by an 850 nm Infrared LED. A 2592 × 1944-pixel CMOS camera (ACA2500-14um, Basler) was used to capture images of the plate at 14 frames per second and a magnification of 20 μm per pixel. Optogenetic light was delivered using three 625 nm LEDs (M625L3, Thorlabs) placed in a way that their light uniformly illuminated the camera’s field of view on the agarose plate. A 3-second light pulse [randomly chosen from either 150 μW/mm^2^ (for stimulation) or 0 μW/mm^2^ (for no stimulation control)] with an inter-trigger interval of 30 seconds was delivered to the agarose plates.

A previously described post-processing algorithm was used to identify worm behaviors through pose dynamics classification, which classified worm pose dynamics into a behavioral map of forward, reverse, and turns.^44,64,65^ To calculate change in velocity, only worms that are moving forward at the stimulus onset were considered. Change in velocity was defined as the difference between the velocity at the end of the 3-second optogenetic stimulus and the velocity at the beginning of the optogenetic stimulus. A threshold of -0.05 mm/s in velocity change was used to define slowdown or reversal. A reversal bout was defined as epochs where instantaneous velocity dropped below a threshold of -0.02 mm/s. The reversal distance was computed as the time integral of absolute velocity during the reversal bout. Reversal duration was calculated as the bout duration (in seconds).

### Microscopy, calcium imaging, and cell ablation

To image synaptic markers, worms were mounted on a 3% agarose pad with various amounts of 2,3-butanedione monoxime (0.75% for L1; 3% for L4; 5% for dauer and adult). To perform calcium imaging and cell ablation, worms were mounted on a 10% agarose pad with 100 nm polystyrene beads (Polysciences #00876–15, mixed with M9 at a 1:5 ratio).

Images were captured on a Leica DMi8 inverted microscope platform equipped with a Leica K5 monochrome camera. Measurements of fluorescent intensity were made using the Leica Application Suite X (3.7.6.25997) software.

For calcium imaging, eggs were placed onto plates seeded with a bacterial lawn containing 5 mM all-trans retinal and then grown in the dark for 3 days at 20°C. Day-one adults were mounted on a 10% agarose pad with 2 μL of 100 nm polystyrene beads (mixed with M9 at a 4:1 ratio). Imaging was done on a Leica DMi8 inverted microscope equipped with an Infinity scanner that has a 638-nm laser. To visualize the calcium transit in AVM neuron following the optogenetic activation of ALM, we used the *unkEx910[T20B3.14p::Chrimson; mec-18p::GCaMP7s::tbb-2 3’ UTR; ceh-22p::GFP]* transgene, which expressed Chrimson in ALM neurons and GCaMP7s in all TRNs. In this application, a 638 nm laser beam (spot size = 0.41 μm; scan speed = 1500; laser power = 1%) was guided to scan the ALMR cell body (“fill” mode) for 3 seconds through a 63x water lens. The GCaMP signal of AVM was then recorded before and after the optogenetic stimulation using blue excitation light and a GFP filter set.

To visualize interneuron activation upon TRN activation, we used the *wtfIs46[mec-4p::Chrimson::SL2::mCherry]* and *sraIs49[nmr-1p::GCaMP3]* transgenes, which expressed Chrimson in all TRNs and GCaMP3 in the interneurons, respectively. Strains all carry a *lite-1(lf)* background. In this application, a 638 nm laser beam (spot size = 0.41 μm; scan speed = 1500; laser power = 100%) was guided to scan the nerve ring (boxed in the “fill” model) for 3 seconds.

To ablate AVM neuron, L2 worms with a TRN marker (*uIs94* or *uIs178*) were mounted on a 10% agarose pad with 1 μL of 100 nm polystyrene beads (mixed with M9 at a 1:5 ratio). A 355 nm UV laser beam (spot size = 0.5 μm; scan speed = 1; laser power = 20%) was guided to scan the AVM cell body (“edge only” mode) through a 63x water lens for one cycle. Laser-treated worms were then rescued using M9 buffer and cultured at 20°C until they reached the L4 or day-one adult stage. The loss of AVM neurons was later confirmed by the absence of red fluorescence signal on the day of the experiment.

### Plate dropping assay

To precisely control the weight of the plate in the plate dropping assay, we designed and custom-built a 3D apparatus that is a plate holder but also contains slots that can be filled with various weights (the blueprint of the device can be found in Data S1). By adjusting the weights added to the plate holder, we can calibrate each experiment, so that the total weight of the plate holder and the NGM plate is always around 350 g. In the plate dropping assay, we also controlled the height of the drop to be 7 cm above the stage of the stereomicroscope used to image the animals. The above setup allows a consistent force to be delivered to the worms on the plate across strains and across experiments.

The movements of worms were recorded before and after the NGM plate was dropped. Each experiment included at least 50 worms, and each experiment was repeated at least three times. Assays were performed and scored in a blinded manner. In response to a drop, animals that switched from moving forward to moving backwards and animals that had backward acceleration were counted as a “backward” response. Animals that switched from moving backward to moving forward or animals that had forward acceleration were counted as a “forward” response. Moving animals that stopped in motion in response to the plate dropping were counted as a “pause”. Animals that did not have any behavioral changes were counted as “no response”.

### P. pacificus biting assay

Wells in a 96-well plate were filled with 150 μL solid NGM one day before the experiment. On the day of the experiment, one *P. pacificus* at the day-one adult stage and one *C. elegans* at the specific stages were transferred to the same well on the 96-well plate. Then, the 96-well plate was stored at 20°C. For the one-hour biting assay, the survival of *C. elegans* was examined after 1 hour. For the 24-hour biting assay, the survival of *C. elegans* was checked at the 1-, 3-, 6-, 9-, and 24-hour mark. Only *P. pacificus* with an Eu mouth form were used in all experiments. At least 20 biological replicates were set up for each assay, and each assay was repeated three times for each condition.

## QUANTIFICATION AND STATISTICAL ANALYSIS

All data (except the high-throughput behavioral studies) were plotted as mean ± SD using GraphPad Prism 8.0. For multiple comparisons, we performed one-way ANOVA with Dunnett’s or Tukey’s correction of the *p* values for all pairs of groups. Student’s *t*-test was used in comparisons between two sets of data. One-sample *t*-test was used to test the significant deviation from 0 for the lateralization index data. Single, double, and triple asterisks indicate *p* < 0.05, *p* < 0.01, and *p* < 0.001, respectively.

For the high throughput optogenetic study, statistical analysis is conducted using stimulus events as the fundamental unit. For each experiment, we reported the proportion of events that result in a certain behavioral outcome, the total number of stimulus events, and the corresponding 95% confidence interval. Kolmogorov-Smirnov test with Bonferroni correction was used to determine the statistical significance for the change in velocity between the stimulation and non-stimulation conditions. Two-proportion Z-tests^66^ with Bonferroni correction were used to analyze the statistical significance when comparing the probability of slowdown or reversals between the stimulation and non-stimulation conditions.

