## Supplementary material for "Synaptic and neural pathway redundancy enables the robustness of a sensory-motor reflex and promotes predation escape in *C. elegans*": Table S1-S3; Figure S1-S7

**Table S1. Innxin expression patterns in the touch circuit.** TPM from the L4 stage scRNA-seq data are shown (Taylor et al., 2021; Threshold 2). The green ones were found to be expressed based on fosmid reporter studies (Bhattacharya et al., 2019).

| | ALM | AVM | PVM | PLM | AVD | PVC |
|---|---|---|---|---|---|---|
| *inx-1* | 149.5 | 94.6 | 78.2 | 196.6 | 103.8 | 67.4 |
| *inx-2* | 0.6 | 0.0 | 0.0 | 0.0 | 0.0 | 0.0 |
| *inx-7* | 15.1 | 14.2 | 1.8 | 19.5 | 35.0 | 60.7 |
| *inx-8* | 0.0 | 5.9 | 0.0 | 0.0 | 0.0 | 1.3 |
| *inx-10* | 10.0 | 6.9 | 0.0 | 4.3 | 4.4 | 14.5 |
| *inx-14* | 0.0 | 0.0 | 0.0 | 0.0 | 2.1 | 0.0 |
| *inx-18* | 0.0 | 0.0 | 0.0 | 0.0 | 0.0 | 0.0 |
| *inx-19* | 0.0 | 0.0 | 0.0 | 0.0 | 0.0 | 84.4 |
| *unc-7* | 45.9 | 25.0 | 23.3 | 129.0 | 55.4 | 84.8 |
| *unc-9* | 69.6 | 241.6 | 101.9 | 135.1 | 200.8 | 114.4 |

**Table S2. Functional screen of innexin genes for their involved in touch response.** The strain is deemed "sensitive" if the animals responded to at least four out of five touches on average (N = 20).

| **Young larva** | | | |
|---|---|---|---|
| Strain name | Genotype | Anterior touch response | Posterior touch response |
| VC40335 | *inx-1(gk580946)* | Sensitive | Sensitive |
| VC260 | *inx-2(ok376)* | Sensitive | Sensitive |
| RB1792 | *inx-7(ok2319)* | Sensitive | Sensitive |
| VC116 | *inx-8(gk42)* | Sensitive | Sensitive |
| RB2051 | *inx-10(ok2714)* | Sensitive | Sensitive |
| AU98 | *inx-14(ag17)* | Sensitive | Sensitive |
| RB1896 | *inx-18(ok2454)* | Sensitive | Sensitive |
| CX6161 | *inx-19(ky634)* | Sensitive | Sensitive |
| CB5 | *unc-7(e5)* | Insensitive | Sensitive |
| CB101 | *unc-9(e101)* | Insensitive | Sensitive |
| **Adults** | | | |
| Strain name | Genotype | Anterior touch response | Posterior touch response |
| VC40335 | *inx-1(gk580946)* | Sensitive | Sensitive |
| VC260 | *inx-2(ok376)* | Sensitive | Sensitive |
| RB1792 | *inx-7(ok2319)* | Sensitive | Sensitive |
| VC116 | *inx-8(gk42)* | Sensitive | Sensitive |
| RB2051 | *inx-10(ok2714)* | Sensitive | Sensitive |

| | | | |
|---|---|---|---|
| AU98 | *inx-14(ag17)* | Sensitive | Sensitive |
| RB1896 | *inx-18(ok2454)* | Sensitive | Sensitive |
| CX6161 | *inx-19(ky634)* | Sensitive | Sensitive |
| CB5 | *unc-7(e5)* | Sensitive | Sensitive |
| CB101 | *unc-9(e101)* | Sensitive | Sensitive |
| CGZ113 | *unc-9(e101); inx-1(gk580946)* | Insensitive | Insensitive |
| CGZ197 | *unc-9(e101); inx-1(unk29)* | Insensitive | Insensitive |
| CGZ72 | *unc-9(e101); inx-2(ok376)* | Sensitive | Sensitive |
| TU4126 | *unc-9(e101); inx-7(ok2319)* | Sensitive | Sensitive |
| CGZ1123 | *unc-9(e101); inx-8(gk42)* | Sensitive | Sensitive |
| CGZ1124 | *unc-9(e101); inx-10(ok2714)* | Sensitive | Sensitive |
| CGZ1125 | *unc-9(e101); inx-14(ag17)* | Sensitive | Sensitive |
| CGZ1126 | *unc-9(e101); inx-18(ok2454)* | Sensitive | Sensitive |
| CGZ1127 | *unc-9(e101); inx-19(ky634)* | Sensitive | Sensitive |
| ZM3087 | *unc-9(fc16); unc-7(e5)* | Sensitive | Sensitive |

**Table S3. Primers used in this study, related to STAR Methods.**

| **Primer name** | **Sequences (5' to 3')** |
|---|---|
| mec-17p_F | ggtgaccattcttcaggtagggtaa |
| mec-17p_R | gatcgaatcgtctcacaactgatcc |
| unc-9_F | atgagtatgctattgtattatttcgcgagtg |
| unc-9_R | cacgtcgtgcatttttccttctttattg |
| ceh-22p_F | tgcatgcaccggatgaagattc |
| ceh-22p_R | cggctctcgagactcgagt |
| nmr-1p_F | gttgtggtctcactttgaacgacaa |
| nmr-1p_R | atctgtaacaaaactaaagtttgtcgtgttcc |
| unc-7_isoform a_F | gtgaacttatgatcctcaaatagaatttttaggacctc |
| unc-7_isoform a_R | gttgtaaaacgacggccagtcggcattccagaaccgaaac |
| inx-1_F | atgcttctatattatctggcggccat |
| inx-1_R | ttagacgaacgtgaagtaaccatctga |
| T20B3.14p_F | cgcaataaactaaagccgacc |
| T20B3.14p_R | gtttctgcaaaatttttgatttttttgtcaaatgaga |
| mec-18p_F | gcttcaattaattcgtctactatccacgtg |
| mec-18p_R | gctcacaaccttcttggaaggc |

| | |
|---|---|
| eat-4(cDNA)_F | atgtcgtcatggaacgaggcttg |
| eat-4(cDNA)_R | ctaccactgctgataatgcggattttc |
| gcy-37p_F | acgttgccgggctaataagtatg |
| gcy-37p_R | atttctgtgtagtagaaaaagtagaaaagcg |
| eat-4_F | atgtcgtcatggaacgaggc |
| eat-4_R | ctaccactgctgataatgcggat |
| eat-4 3'UTR_R | caagagaggaaagttgagca |
| unc-119(Cb gDNA)_F | cgcccttctattacagggtttctg |
| unc-119(Cb gDNA)_R | gacattctctaatgaaaaaatctttcagttgaaattga |
| ChR2_eYFP_F | atggattatggaggcgccctg |
| ChR2_eYFP_R | ttacttgtacagctcgtccatgc |
| syd-2_F | atgagctacagcaatggaaacataaattgtg |
| syd-2_R | ctaggtatataaatgaaactcgtaggattttgctatgg |
| unc-24_F | atgacgttctcttcttcgtacaaatattcg |
| unc-24_R | ttaaagccaatctgacattcgctcc |
| C10C5.7p_F | tcaacaacttgggcaatgtc |
| C10C5.7p_R | ctgaaattctagaaagatttcgaaagaaaatgca |
| unc-9(gDNA)_F | cgtcgtactttatgtgctcgtgc |
| unc-9(gDNA)_R | caggaaacagctatgaccatatggcaaagtttgtgggttcc |
| CRISPR-inx-1-exon1 | ttctaatacgactcactatagataatatagaagcattccgggttttagagctaga |
| CRISPR-inx-1-exon3 | ttctaatacgactcactatagtgtggagaggactgctgcacgttttagagctaga |
| CRISPR_glc-1_up | ttctaatacgactcactatagcaatggctacctggattgtgttttagagctaga |
| CRISPR_glc-1_down | ttctaatacgactcactataggacagttccactgaatcgcgttttagagctaga |
| CRISPR_glc-2_up | ttctaatacgactcactatagatgtgatatggaggaagagggttttagagctaga |
| CRISPR_glc-2_down | ttctaatacgactcactatagggactgtgagaaatgccgggttttagagctaga |
| CRISPR_glc-3_up | ttctaatacgactcactatagatgtgacctccagagacgggttttagagctaga |
| CRISPR_glc-3_down | ttctaatacgactcactatagctgttccgcaatcataaccagttttagagctaga |
| CRISPR_eat-4_exon2 | ttctaatacgactcactatagatacacttacatttgctgtggttttagagctaga |
| CRISPR_eat-4_exon4 | ttctaatacgactcactatagaatagatgagcttagtgtcagttttagagctaga |
| CRISPR_unc-9_up | ttctaatacgactcactatagctgtacaccgtgaacatcgtgttttagagctaga |
| CRISPR_unc-9_down | ttctaatacgactcactatagaatcgagggatataaaatgggttttagagctaga |

**Supplemental Figures S1-S7**

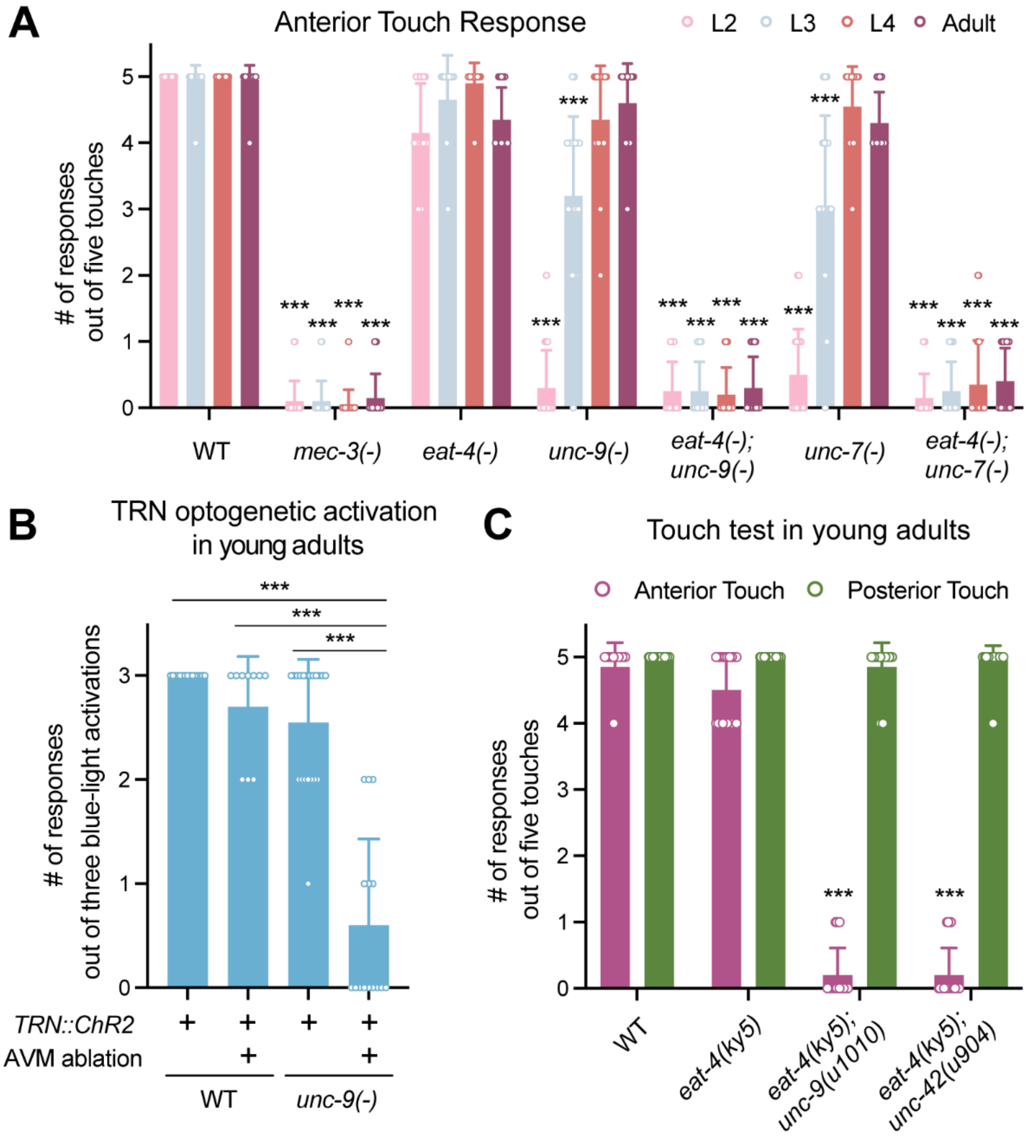


**Figure S1. Anterior touch sensitivity is redundantly controlled by UNC-9::UNC-7 gap junction and glutamatergic synapses, related to Figure 1.** (A) The number of responses out of five anterior touches (by an eyebrow hair) of wild-type, *mec-3(u184)*, *eat-4(ky5)*, *unc-9(e101)*, *eat-4(ky5); unc-9(e101)*, *unc-7(e5)*, and *eat-4(ky5); unc-7(e5)* animals at different developmental stages. Three asterisks indicate $p < 0.001$ in comparison with the wild-type animals at the same stage in a post-ANOVA Dunnett's test. N = 20 for each strain. (B) The number of responses out of three anterior blue-light stimuli in strains carrying the *uIs94[mec-17p::ChR2::YFP; myo-3p::mCherry]* transgene after AVM ablation. N ≥ 10 for all conditions. (C) The number of responses out of five anterior or posterior touches in the indicated strains. *u1010* and *u904* were isolated from a forward genetic screen using *eat-4(ky5)* as the starter strain. N = 20. Mean ± SD is shown.

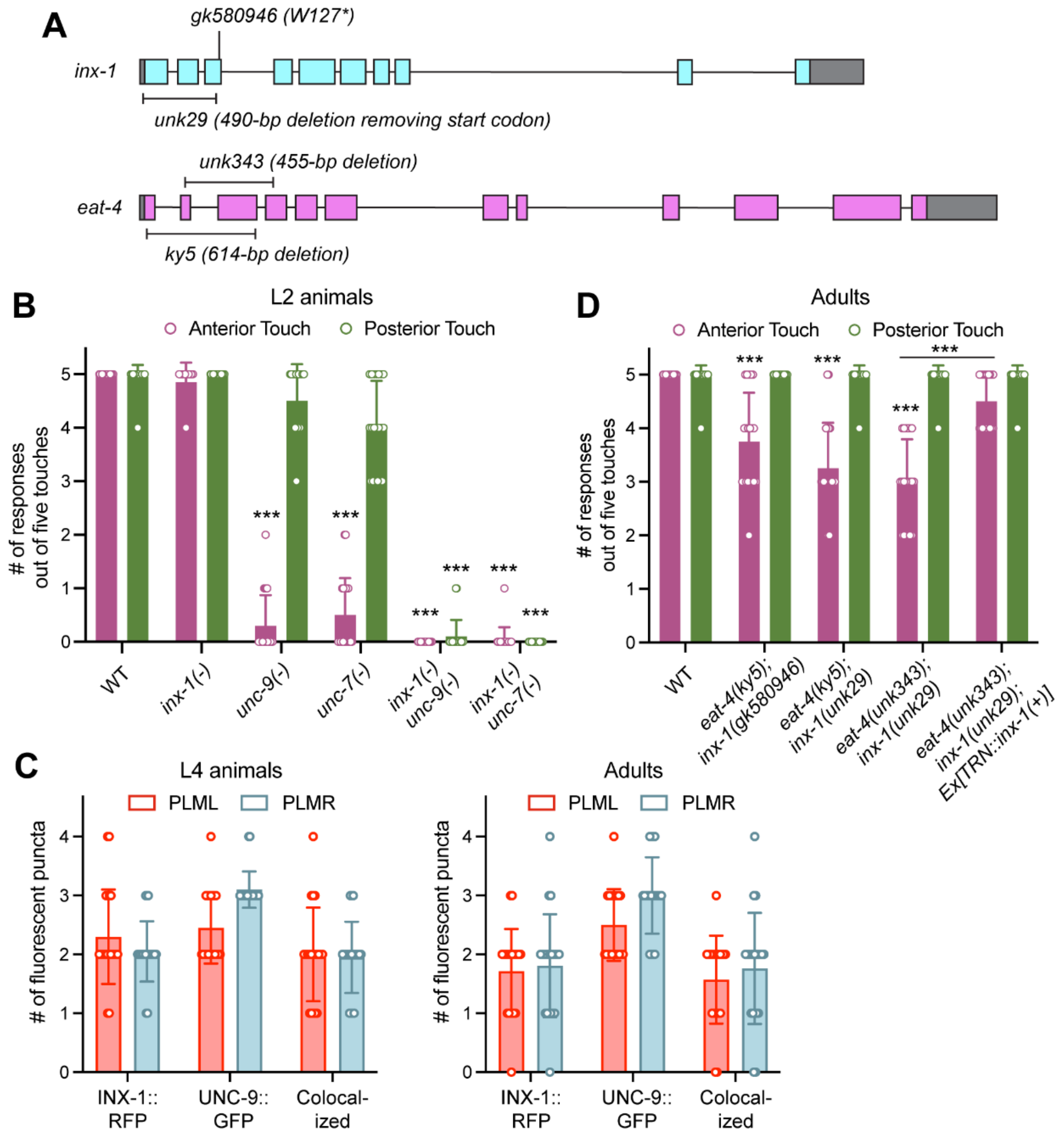


**Figure S2. INX-1 contributes to both anterior and posterior touch sensitivity, related to Figure 2.** (A) Molecular lesions in *inx-1(gk580946)*, *inx-1(unk29)*, *eat-4(ky5)*, and *eat-4(unk343)* alleles; *unk29* and *unk343* are deletion alleles created in this study by CRISPR gene editing. (B) The number of responses out of five anterior or posterior touches in the wild-type, *inx-1(unk29)*, *unc-9(e101)*, *unc-7(e5)*, *inx-1(unk29) unc-9(e101)*, and *inx-1(unk29) unc-7(e5)* animals at the L2 stage. Three asterisks indicate $p < 0.001$ in a post-ANOVA Dunnett's test comparing the mutants with the wild-type animals. N = 20. (C) The number of fluorescent puncta for INX-1::RFP and UNC-9::GFP and the ones with both RFP and GFP signals in the PLML/R anterior neurite segment close to the cell body at both L4 and adult stages. N = 20. Mean ± SD is shown in this Figure. (D) Touch response of the indicated strains; *unkEx769[mec-17p::inx-1a]* was used for TRN-specific rescue of *inx-1*. Three asterisks indicate p < 0.001 in a Tukey's tests comparing the mutants with the wild-type animals between selected pairs. N = 20.

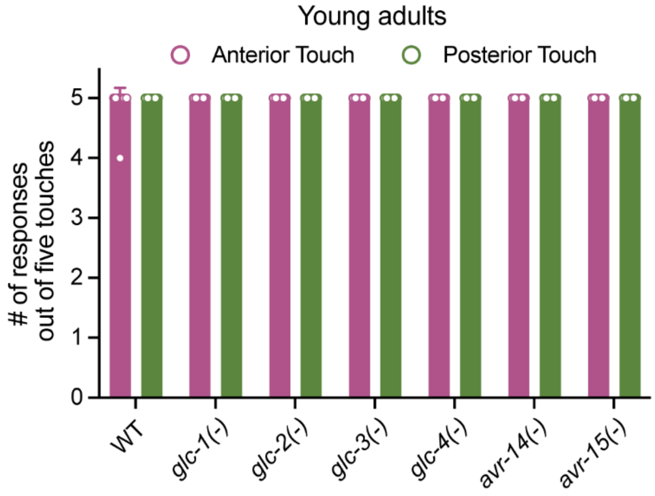


**Figure S3. *glc* single mutants are touch sensitive, related to Figure 3.** The number of responses (mean ± SD) out of five anterior or posterior touches of wild-type, *glc-1(u1053)*, *glc-2(ok1047)*, *glc-3(ok321)*, *glc-4(ok212)*, *avr-14(ad1302)*, and *avr-15(ad1051)* animals. The *u1053* allele is a deletion allele created in this study by CRISPR gene editing. N = 20 for each strain.

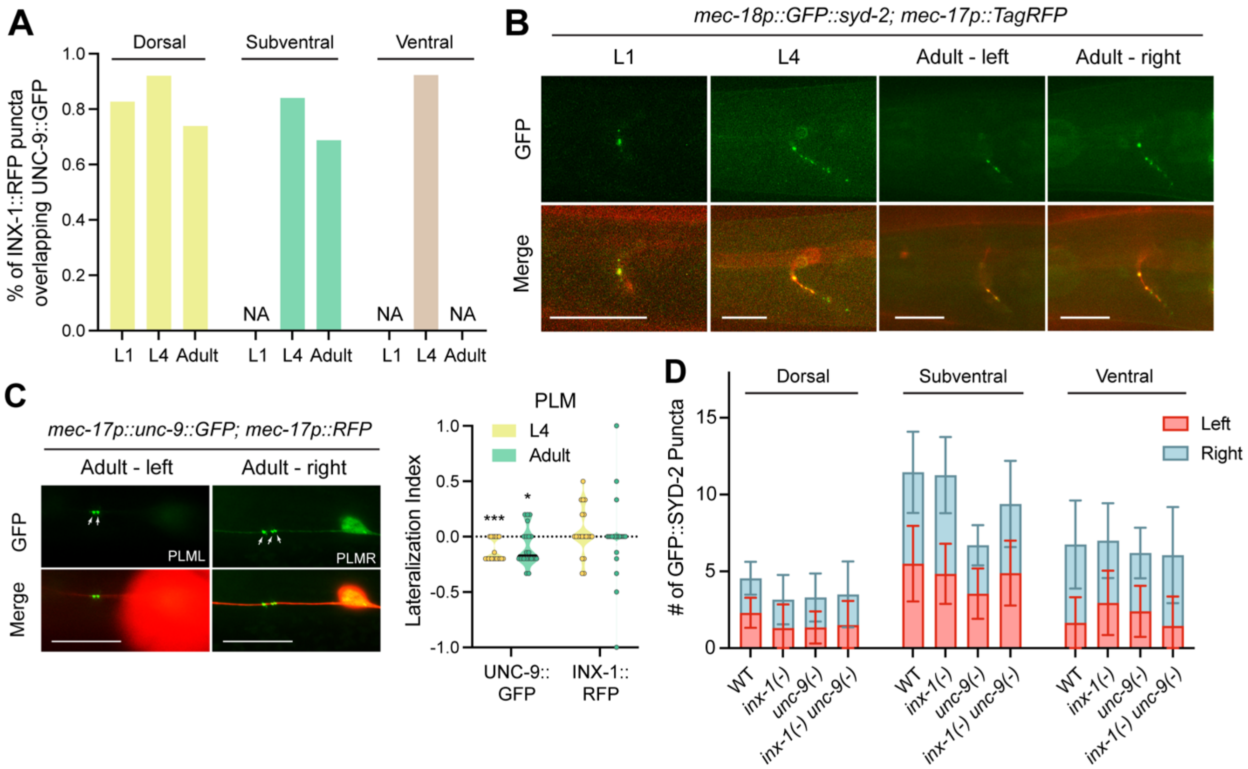


**Figure S4. Localization of synaptic markers in the touch circuit, related to Figure 4.** (A) The percentage of INX-1::RFP puncta that colocalize with UNC-9::GFP puncta in the dorsal, subventral, and ventral zones. Since each animal has only a few INX-1::RFP dots, data from 20 animals were pooled to generate the plot. “NA” means “Not applicable” because there were in total fewer than 5 INX-1::RFP puncta in the pooled data. (B) Representative images of the chemical synapse marker GFP::SYD-2 expressed from the TRN-specific *mec-18* promoter. Scale bars = 20 µm. (C) Representative images of the UNC-9::GFP puncta in PLML and PLMR. Arrows indicate the spatially resolvable puncta. Scale bars = 20 µm. Violin plot of the lateralization index, defined as (# of puncta on the left side - # of puncta on the right side)/total # of puncta from both sides, for UNC-9::GFP and INX-1::RFP in the PLM anterior neurite. One and three asterisks indicate $p < 0.05$ and 0.001, respectively, in a one-sample $t$-test for deviation from 0. (D) Number of GFP::SYD-2 puncta in the various zones of wild-type, *inx-1(unk29)*, *unc-9(e101)*, and *inx-1(unk29); unc-9(e101)* animals. Puncta from the left and right sides were labeled with different colors in the stacked bars, representing the mean ± SD of 20 animals.

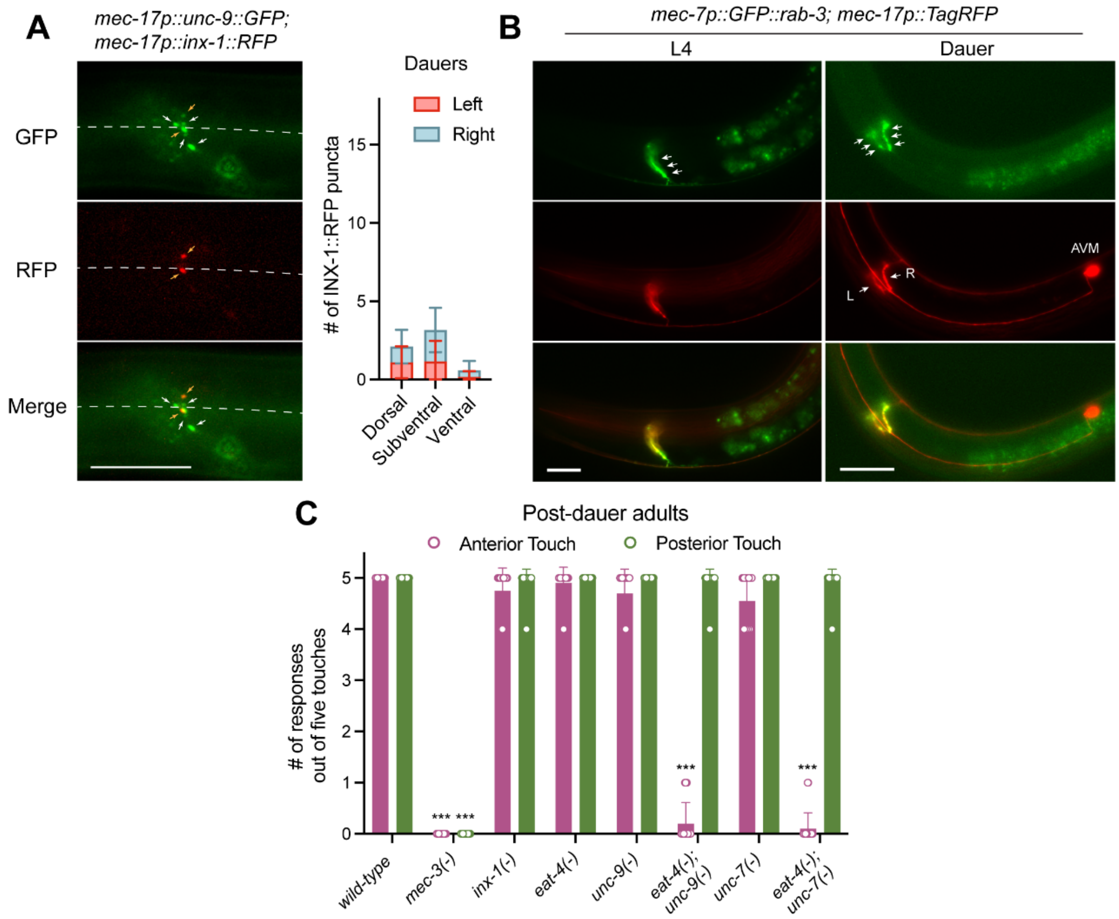


**Figure S5. TRN synaptic markers in dauers, related to Figure 4.** (A) Representative images of UNC-9::GFP and INX-1::RFP in the TRNs of dauers. Orange arrows indicate puncta that are labeled by both GFP and RFP, while white arrows indicate GFP-only puncta. The right panel shows the number (mean ± SD) of INX-1::RFP puncta in dauers. N = 20. (B) Representative images showing the localization of synaptic vesicle markers GFP::RAB-3 in the synaptic branches of AVM in normal L4 and dauer animals. Arrows indicate the GFP signal for RAB-3 at the synapse. AVM in dauers had already grown both left (L) and right (R) synaptic branches that grow into the nerve ring. Scale bars = 20 µm. (C) The number of responses (mean ± SD) out of five anterior or posterior touches in the post-dauer adults of the indicated strains. *mec-3(u184)*, *inx-1(unk29)*, *eat-4(ky5)*, *unc-9(e101)*, and *unc-7(e5)* alleles were used. Three asterisks indicate $p < 0.001$ in a post-ANOVA Dunnett's test comparing the mutants with the wild-type animals. N = 20.

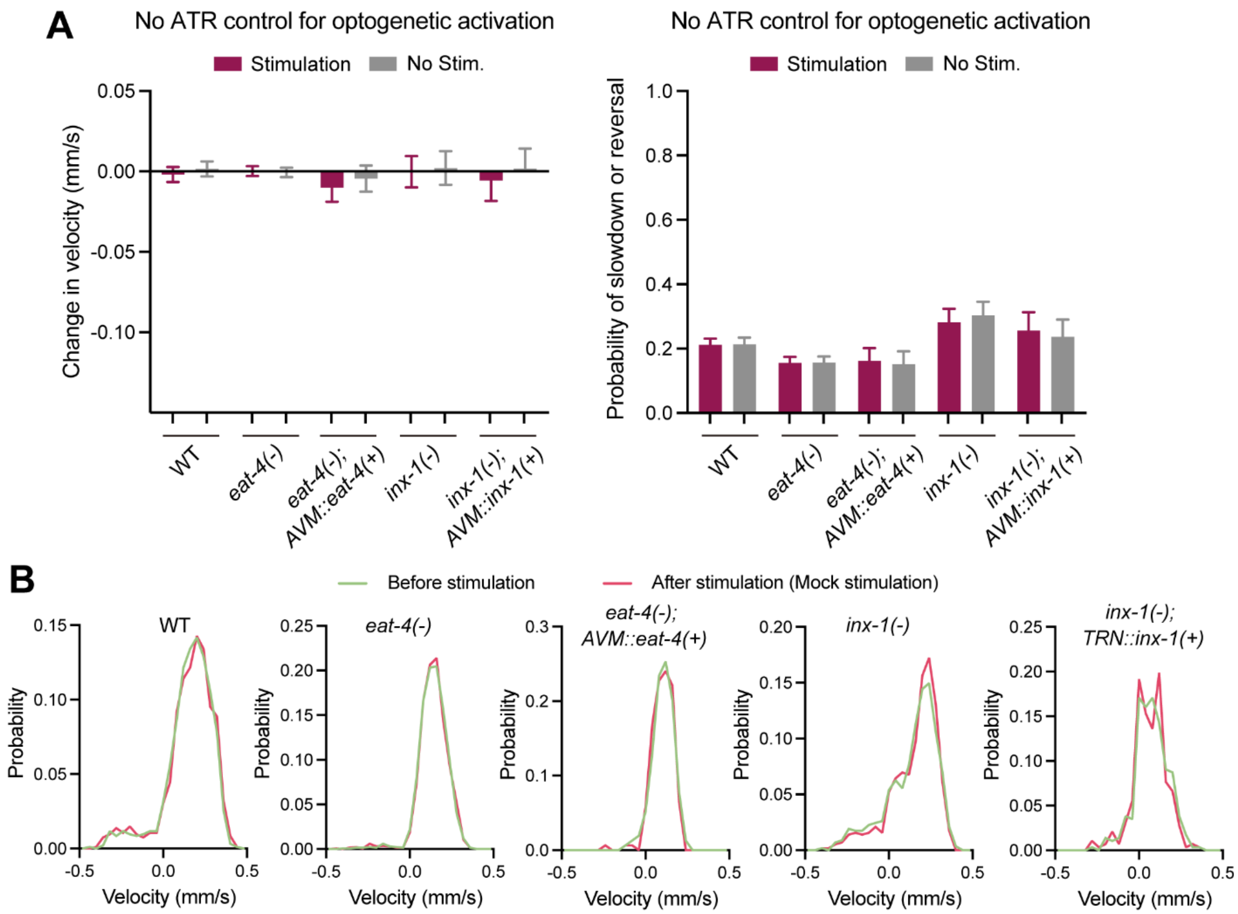


**Figure S6. Control experiments for optogenetic activation of TRNs and high-throughput behavioral analysis, related to Figure 6.** (A) Changes in velocity and probability of slowdown/reversal upon light stimulation (and no stimulation controls) in various strains grown on plates without *all-trans* retinal (ATR). CGZ2400 *wtfIs458[mec-4p::Chrimson]* (N > 1700), CGZ2399 *eat-4(ky5); wtfIs458* (N > 1500), CGZ2529 *eat-4(ky5) III; wtfIs458; unkEx765[gcy-37p::eat-4]* (N > 350), CGZ2519 *inx-1(unk29); wtfIs458* (N > 550), and CGZ2568 *inx-1(unk29); wtfIs458; unkEx769[mec-17p::inx-1]* (N > 300) were used. Mean ± 95% confidence interval is shown. No significant difference was found between stimulation and no stimulation conditions in the Kolmogorov-Smirnov test for the left panel and the two-proportion Z-test for the right panel. (B) Velocity distribution before and after the mock stimulation (no stimulation) in the presence of ATR. No statistical significance in the Kolmogorov-Smirnov tests was found.

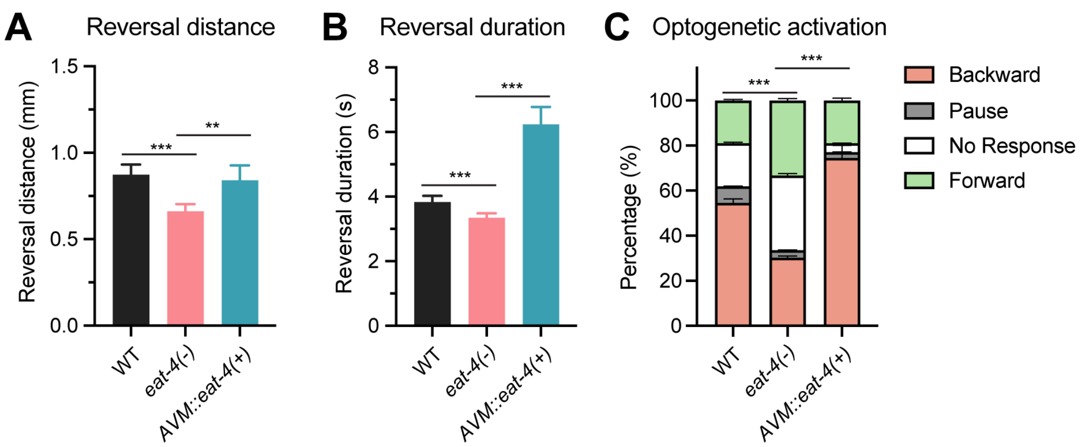


**Figure S7. AVM expression of *eat-4* contributes to reversal behavior upon optogenetic TRN activation, related to Figure 7.** (A) Comparison of reversal distance in CGZ2400 *wtfIs458[mec-4p::Chrimson]* (N = 367), CGZ2399 *eat-4(ky5); wtfIs458* (N = 313), and CGZ2529 *eat-4(ky5) III; wtfIs458; unkEx765[gcy-37p::eat-4]* (N = 83) upon red light activation of TRNs. (B) Comparison of reversal duration in the three strains. Three and two asterisks indicate $p < 0.001$ and 0.01, respectively, in Tukey's multiple comparisons tests. (C) Percentage of behavioral response of the three strains upon red light stimulation. Three asterisks indicate $p < 0.001$ in a two-proportion Z test considering only the "backward" response. Mean ± 95% confidence interval is shown in this Figure.
