## Supplementary material for "Synaptic and neural pathway redundancy enables the robustness of a sensory-motor reflex and promotes predation escape in *C. elegans*": Data S1

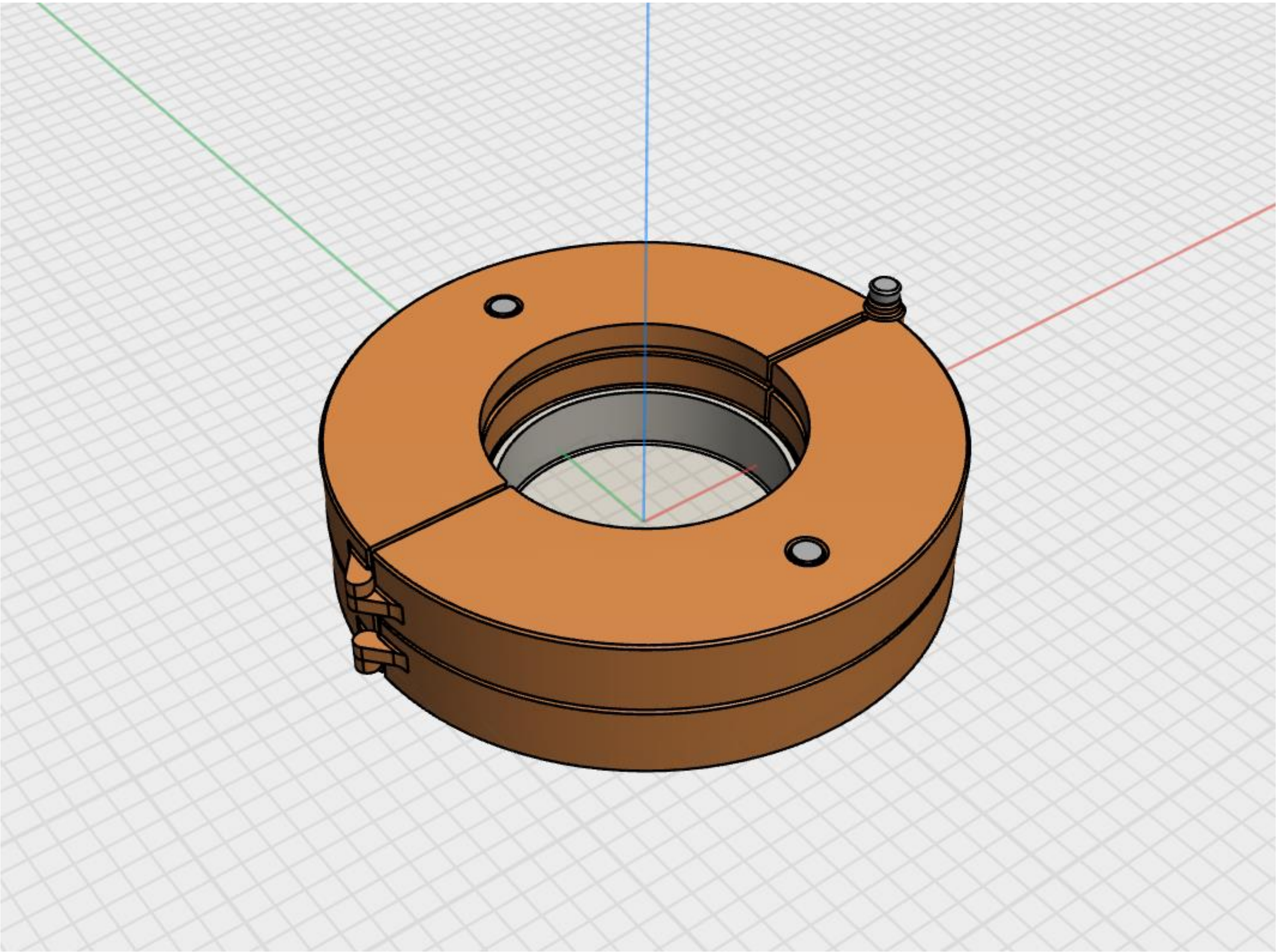

This is the Plata Dropping Apparatus designed for this study. The whole Apparatus is a clamp consisting of two arms. When the two arms close, they can tightly clamp a 60-mm NMG plate.

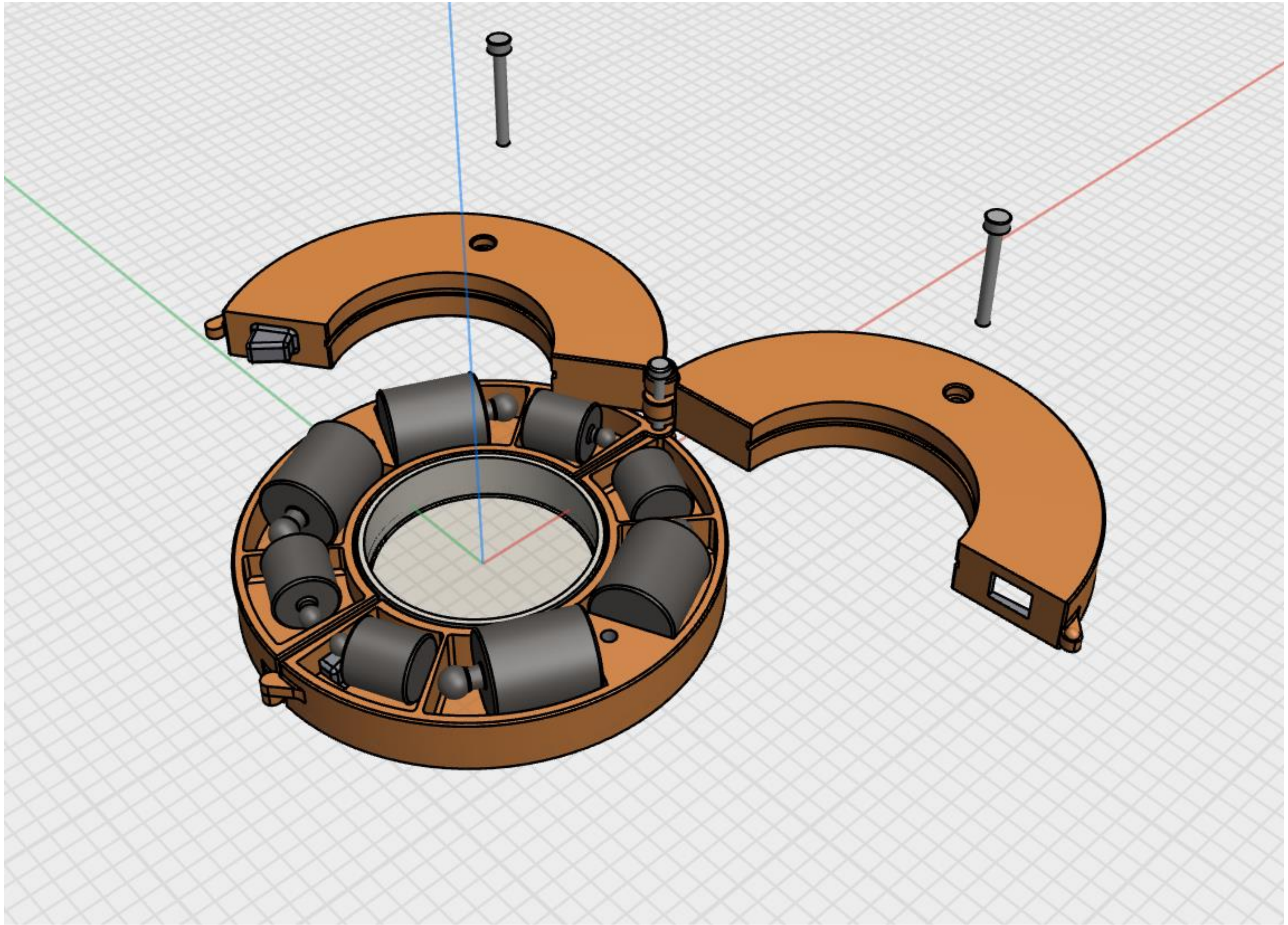

This Figure shows the inside of the Apparatus. There are four cavities in each arm; two are for the 50-gram weight and the other two are for the 2-gram weight. Before experiments, desired weights were put into the Apparatus and crannies were filled with clay to fix the weights.

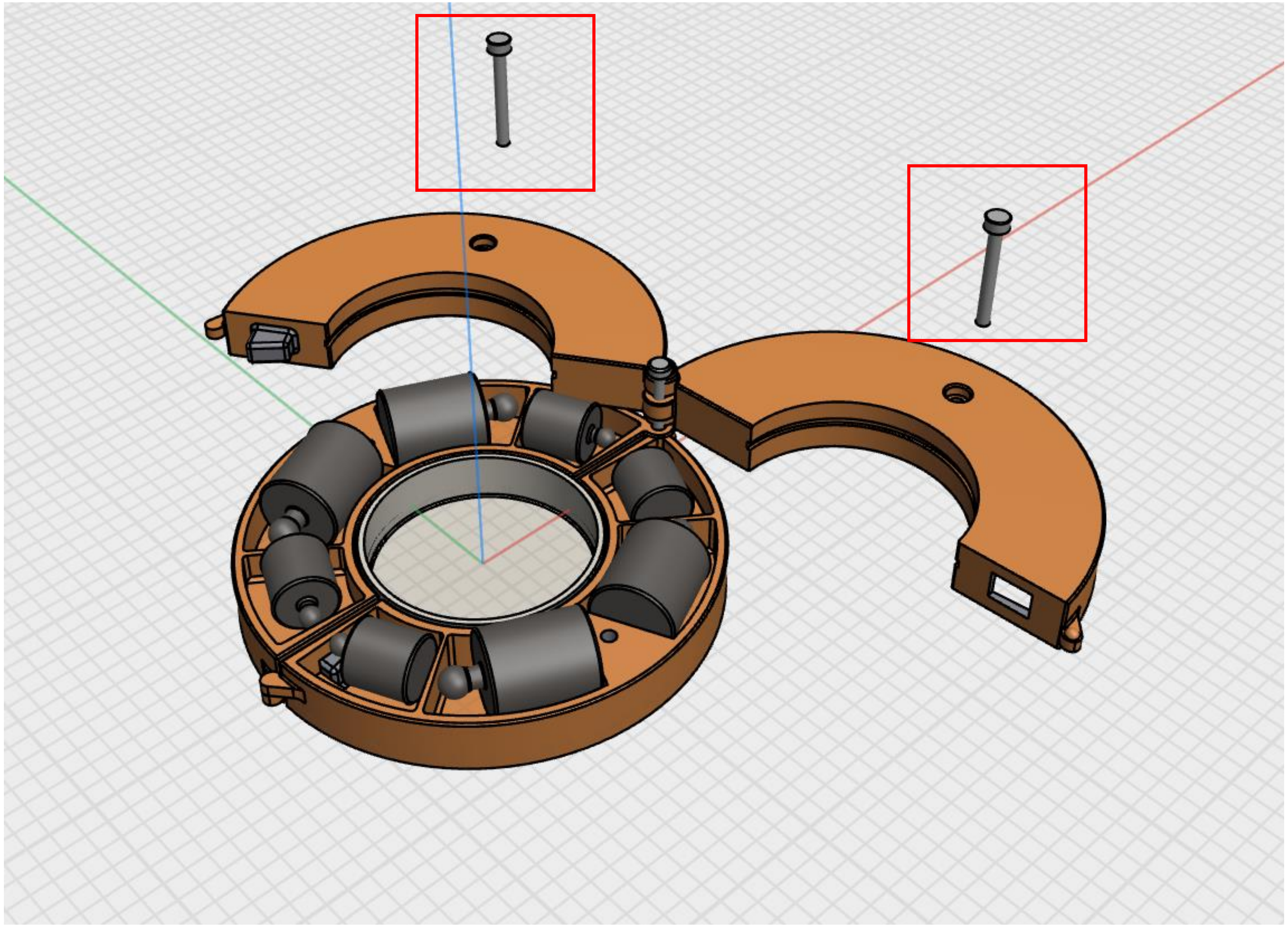

Two bolts marked by red boxes are designed to lock the arms.
